# Structural plasticity of bacterial ESCRT-III protein PspA in higher-order assemblies

**DOI:** 10.1101/2024.07.08.602472

**Authors:** Benedikt Junglas, Esther Hudina, Philipp Schönnenbeck, Ilona Ritter, Anja Heddier, Beatrix Santiago-Schübel, Pitter F. Huesgen, Dirk Schneider, Carsten Sachse

## Abstract

Eukaryotic members of the *endosome sorting complex required for transport* III (ESCRT-III) family have been shown to form diverse oligomeric assemblies. The bacterial *phage shock protein A* (PspA) has recently been identified as a bacterial member of the ESCRT-III superfamily, and monomeric PspA homo-oligomerizes to form large rod-shaped assemblies. As observed for eukaryotic ESCRT-III, PspA forms different tubular assemblies with varying diameters. Using electron cryo-microscopy (cryo-EM), we determined a total of 61 PspA structures and observed in molecular detail how structural plasticity of PspA rods is mediated by conformational changes at three hinge regions in the monomer and by the fixed as well as changing molecular contacts between protomers. Moreover, we reduced and increased the structural plasticity of PspA rods by removing the loop connecting helices α3/α4 and the addition of nucleotides, respectively. Based on our analysis of PspA-mediated membrane remodeling, we suggest that the observed mode of structural plasticity is a prerequisite for the biological function of ESCRT-III superfamily members.

**Summary:** A series of cryo-EM structures of PspA rods with induced diameter modulations reveals the molecular basis of structural plasticity.

## Introduction

In eukaryotes, ESCRT (Endosomal sorting complexes required for transport) proteins form a multi-subunit machinery that performs a topologically unique membrane budding reaction away from the cytoplasm (Schmidt and Teis, 2012). The ESCRT system assumes critical roles in many cellular processes, including nuclear envelope sealing (Vietri et al., 2015), plasma membrane repair, lysosomal protein degradation (Zhu et al., 2017), retroviral budding, and the multivesicular body (MVB) pathway (Schmidt and Teis, 2012). In particular, the conserved group of ESCRT-III proteins is associated with such membrane remodeling processes, and their biological activity appears to be linked to the formation of homo and/or heteropolymeric structures, such as sheets, strings, rings, filaments, tubules, domes, coils, and spirals (Huber et al., 2020; McCullough et al., 2015; Pfitzner et al., 2021, 2020). Based on cryo-EM structures, it was recently shown that the bacterial proteins *phage shock protein A* (PspA) and the vesicle-inducing protein in plastids 1 (Vipp1, also known as *the inner membrane-associated protein of 30 kDa*, IM30) are structurally similar to ESCRT-III proteins found in eukaryotes and archaea (Gupta et al., 2021; Junglas et al., 2021; Liu et al., 2021b). Simultaneously, it has been demonstrated by phylogenetic analyses that the ESCRT-III and PspA proteins indeed have a common ancestor, thereby extending the ESCRT-III superfamily to the bacterial domain (Di Giulio, 2021; Liu et al., 2021b).

The 25.3 kDa PspA is the main effector of the *E. coli* Psp system and is thought to protect and stabilize the bacterial inner membrane by a so far unknown mechanism (Kleerebezem and Tommassen, 1993; Kobayashi et al., 2007). In bacteria, PspA is present both in a soluble form in the cytoplasm and bound to negatively charged lipids of the inner membrane (Hankamer et al., 2004; Jovanovic et al., 2014). Thereby, PspA can counteract membrane stress and, for example, block the leakage of protons through damaged membranes (Joly et al., 2010; Kobayashi et al., 2007; Manganelli and Gennaro, 2017). Another well-studied example of a PspA-like protein that plays an essential role in membrane protection but also in membrane remodeling (*i.e.,* of the thylakoid membrane) is the PspA homolog Vipp1. Vipp1 is capable of membrane fusion as a means of active membrane repair, as well as passive membrane protection of a protein carpet forming on the membrane (Hennig et al., 2015; Junglas et al., 2020a; Siebenaller et al., 2019). PspA and Vipp1 appear to share a common ancestor and share structural similarities. Both proteins solely consist of α-helices connected by short loops (Vipp1 seven α-helices, PspA six α-helices), with a four helix-core structure, in which helix α1 and helix α2/3 form a coiled-coil hairpin connected by a short loop (Gupta et al., 2021; Junglas et al., 2021; Liu et al., 2021b). A distinctive feature of PspA and Vipp1 is their ability to form MDa-sized ring or rod-shaped homo-oligomers, as observed also with eukaryotic ESCRT-IIIs (Gupta et al., 2021; Junglas et al., 2021; Liu et al., 2021b; Male et al., 2014; Theis et al., 2019). PspA rods are helical assemblies with the monomers arranged by a brick-like stacking, while the four-helix core of helix α1-α4 forms the wall of the tubular PspA rods, and the amphipathic helix α5 pointing to the outside (Junglas et al., 2021). Recently the molecular structure of a rod with a diameter of about 200 Å was solved by cryo-EM, PspA of the cyanobacterium *Synechocystis* sp. PCC 6803 (from here on PspA) was shown to form diverse rods having variable diameters and lengths of up to several hundred nanometers (Junglas et al., 2021).

Eukaryotic ESCRT-III monomers have been observed in different conformational states in formed heteropolymeric complexes of different ESCRT-III isoforms (Azad et al., 2023; Nguyen et al., 2020). However, the structural flexibility of ESCRT-III proteins in homopolymers and the enabling structural mechanisms have not been studied in molecular detail. We now solved the structures of 61 PspA rods using cryo-EM and observed a wide range of finely sampled assemblies with different rod diameters displaying a notable structural plasticity. We show that nucleotide addition affects the distribution of rod diameters. In the absence of nucleotides, PspA rods have narrow diameters, PspA rods shift to wider diameters in the presence of ADP until in the presence of ATP, PspA rods have the widest diameters. Notably, we show that the addition of ATP enhances PspA’s membrane remodeling activity indicating that structural plasticity might be critical for PspA’s ability to engulf and remodel membranes. Comparing the structures of 11 individual PspA structures within varying diameters, we observed that the structural plasticity of PspA rods requires conformational changes at three defined hinge regions. In these structures, the length of helix α3 increases as the loop connecting helices α3 and α4 shortens within larger diameter rods. Upon deletion of the loop connecting helices α3/α4, we observed a significant reduction of the structural plasticity and no effect on the diameter distribution upon addition of ATP. In summary, PspA monomers display a remarkable structural plasticity allowing the assembly of multiple stable rod structures with varying diameters.

## Results

### PspA forms a series of helical rod-shaped assemblies with different diameters

To investigate the structural heterogeneity observed in previous PspA analyses (Junglas et al., 2021), we prepared PspA rods by refolding PspA purified in denaturing conditions, and observed the unassisted formation of rods with variable diameters ranging from 180 to 250 Å with a maximum of the distribution at 215 Å (**Fig. 1A**). After multiple rounds of classification and symmetry analysis of helical segments (**Fig. 1B**), we determined the cryo-EM structures of rods with 180, 200, 215, 235 and 250 Å in diameter between 4.2 and 6.2 Å resolution. The 200 Å rod structure determined at 4.2 Å resolution is very similar to the previously published structure of PspA rods (PDB:7ABK, EMDB: 11698), whereas increasing diameters showed lower resolution (**Fig. 1C**). The cross-sectional top and side views of the cryo-EM maps show features expected for PspA rods (spikes formed by the α1/2+3 hairpin, Christmas-tree pattern in the cross-sectional side view). The continuous series of PspA cryo-EM structures confirms that PspA on its own is capable of forming a range of different stable assemblies indicating structural plasticity.

**Figure 1:**
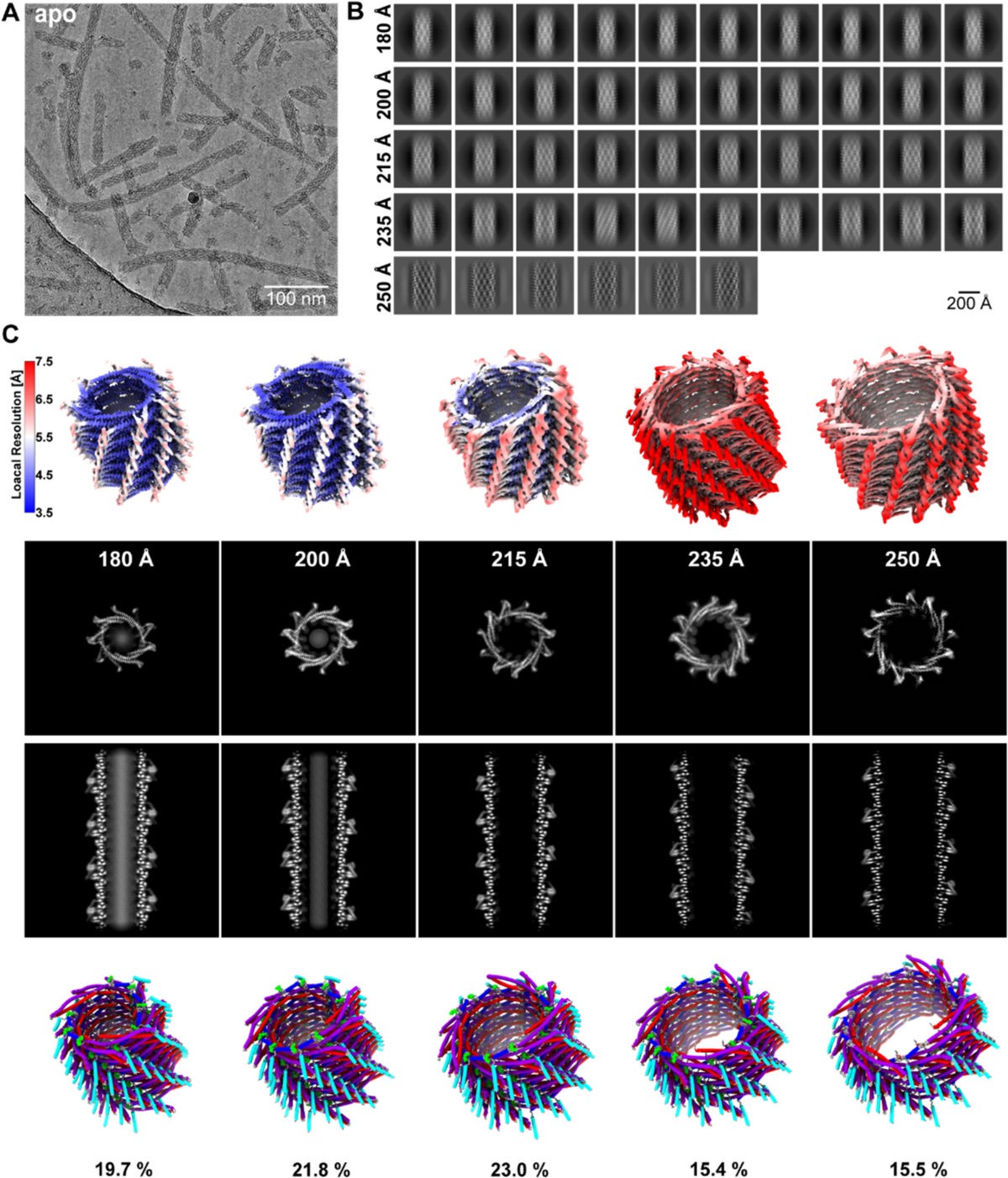
PspA self-assembles into a series of different rods with diameters from 180 to 250 Å. **A:** Cryo-EM micrograph of PspA revealing different rods with respect to the length and width of the observed assemblies. **B:** Image classification of helical segments into five different groups with diameters of 180, 200, 215, 235, and 250 Å, respectively. **C:** Five corresponding three-dimensional cryo-EM structures colored with local resolution estimates (global resolution 4.3, 4.2, 4.6, 5.7, 6.2 Å, respectively). Cross-sectional top and side view (top) of the five structures with increasing diameters including the atomic models (bottom) with the respective relative abundance, with helices colored as follows: α1 red, α2+3 violet, α4 blue, α5 cyan.

### Structural plasticity of PspA rods can be enhanced by the addition of ADP and ATP

Recently, the PspA-related protein Vipp1 was shown to form tapered ring structures with different conformations and diameters inside the determined assembly (Gupta et al., 2021). Moreover, as nucleotides were identified in the narrowest ring, it had been suggested that ADP and/or ATP binding affected the diameter of the assembly. Inspired by this observation, we added ADP during filament formation and measured a diameter distribution broadened to a range from 180 to >400 Å with a maximum at 250 Å, whereas the fraction of rods having a diameter > 280 Å did not exceed 5% (Fig. 2A). Acquiring cryo-EM data for the ADP sample, we determined a total of 11 additional cryo-EM structures (at 4.3 to 7.6 Å global resolution) for the respective diameters. A total of five structures (180, 200, 215, 235, and 250 Å) out of the 11 structures completely overlapped with the structures solved above in the absence of nucleotides. However, seven additional rod structures with increased diameters were solved and, interestingly, rods with diameters >320 Å contained additional cylindrical density in the lumen. The corresponding micrographs suggested that “super rods” were formed by large rods engulfing smaller PspA rods of different helical symmetry and appeared as smooth cylinders after the imposition of helical symmetry (Suppl. Fig. 1A). Finally, we also incubated PspA with ATP during tube formation and were able to determine a total of 11 additional cryo-EM structures at resolutions ranging from 3.8 to 6.9 Å (Fig. 2B). It is important to note that for rod diameters 180, 200, 215 and 235 Å we identified density in the putative nucleotide binding site close to the loop connecting helices *α*3 and *α*4 (aa 155 – 156), albeit weaker than the surrounding protein density presumably due to incomplete binding (Suppl. Fig. 1B). Closer inspection of the binding pocket and comparison with the PspA apo model revealed that the nucleotide competes with the α3/α4 loop, which results in an alternative conformation and reduced mobility of the α3/α4 loop, associated with lower atomic temperature factors, when the nucleotide is present (Suppl. Fig. 1C). While some of the rod structures were already identified in the absence of nucleotides and/or the presence of ADP, the structures of three additional diameters at 290, 305 and 320 Å could be resolved solely when ATP was present. As they represent a large range of structures observed under the same experimental condition at a suitable resolution (**Suppl. Fig. 2A-F**), we built the corresponding 11 atomic models (Fig. 2C). In total, we determined 27 structures of PspA assemblies using cryo-EM in three conditions of nucleotide absence, ADP and ATP, with 14 unique structures differing in their respective diameter, thus enhancing the observed structural plasticity of PspA assemblies.

**Figure 2:**
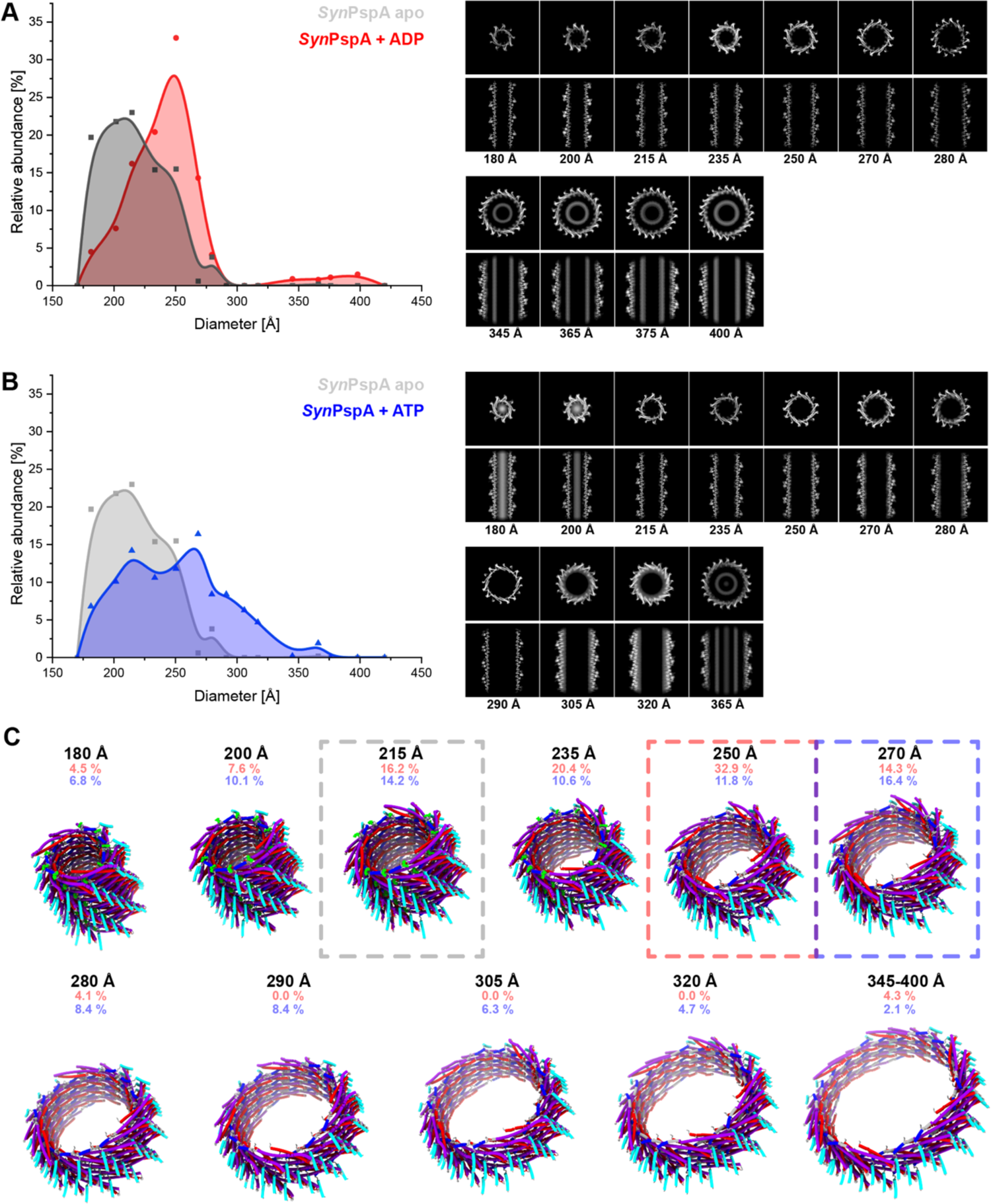
The structural plasticity of PspA rods increases upon incubation with ADP and ATP. **A:** Histogram of rod diameters and the normalized abundance observed upon addition of ADP (red) during refolding in comparison with PspA apo (grey). In the presence of ADP, the distribution maximum shifts to 250 Å and diameters up to 400 Å could be observed. The associated cryo-EM structures are shown in the cross-sectional top and side views. **B:** Histogram of rod diameters upon addition of ATP (blue) during refolding in comparison with PspA apo (grey). In the presence of ATP, the rod population shifts further towards larger diameters in both cases. **C:** Corresponding three-dimensional cryo-EM atomic models. Frame highlights the most frequent structure for *apo* (215 Å, grey), ADP (250Å, red), and ATP (270 Å, blue) (α1 red, α2+3 violet, α4 blue, α5 cyan).

### Three hinge regions and flexible contacts represent the structural dials for PspA’s structural plasticity

To further analyze the cryo-EM maps and elucidate the detailed molecular mechanism underlying the structural plasticity of PspA, we scrutinized the 11 built atomic models. The basic monomer structures were overall similar or nearly identical to the published structure of PspA (PDB:7ABK) in the case of 200 Å diameter rods, except for deviations in the loop connecting helices α3 and α4 (Fig. 3A, Suppl. Fig. 2G). After superimposition of PspA monomer structures from different diameters, we found that five helices α1-α5 were almost unchanged despite changes in their relative orientation (Fig. 3B**, Suppl.** Fig. 3A). In addition, we observed that the helices α3 and α4 become extended with increasing rod diameters resulting in shortening and ultimate disappearance of the loop between these helices in the widest rods (320 and 365 Å diameter). Thus, helices α3 and α4 eventually fused to form a single α-helix. Notably, the length of helix α5 and the loop connecting helices α4 and α5 remained unaffected. The measured atomic distances from Cα-G82 to Cα-P187 increased for PspA rods with smaller diameters (180 – 270 Å), while they decreased again for diameters larger than 270 Å (Fig. 3C). Thus, the PspA molecule adopted a bow-like shape under tension for the smaller diameters, up to an almost linear shape at around 270 Å diameter and a slightly inverted bow curvature for the largest diameters. In more detail, the structural changes were realized by three hinge regions in the monomer: the first hinge was positioned at the transition between α2 and α3 (aa 128-133), the second hinge was located in the loop region between α3 and α4, and third hinge is in the loop region between α4 and α5. The largest angular changes came from the second hinge (−45° to 20°), while the angular changes of the other two hinges were shallower (Fig. 3D).

**Figure 3:**
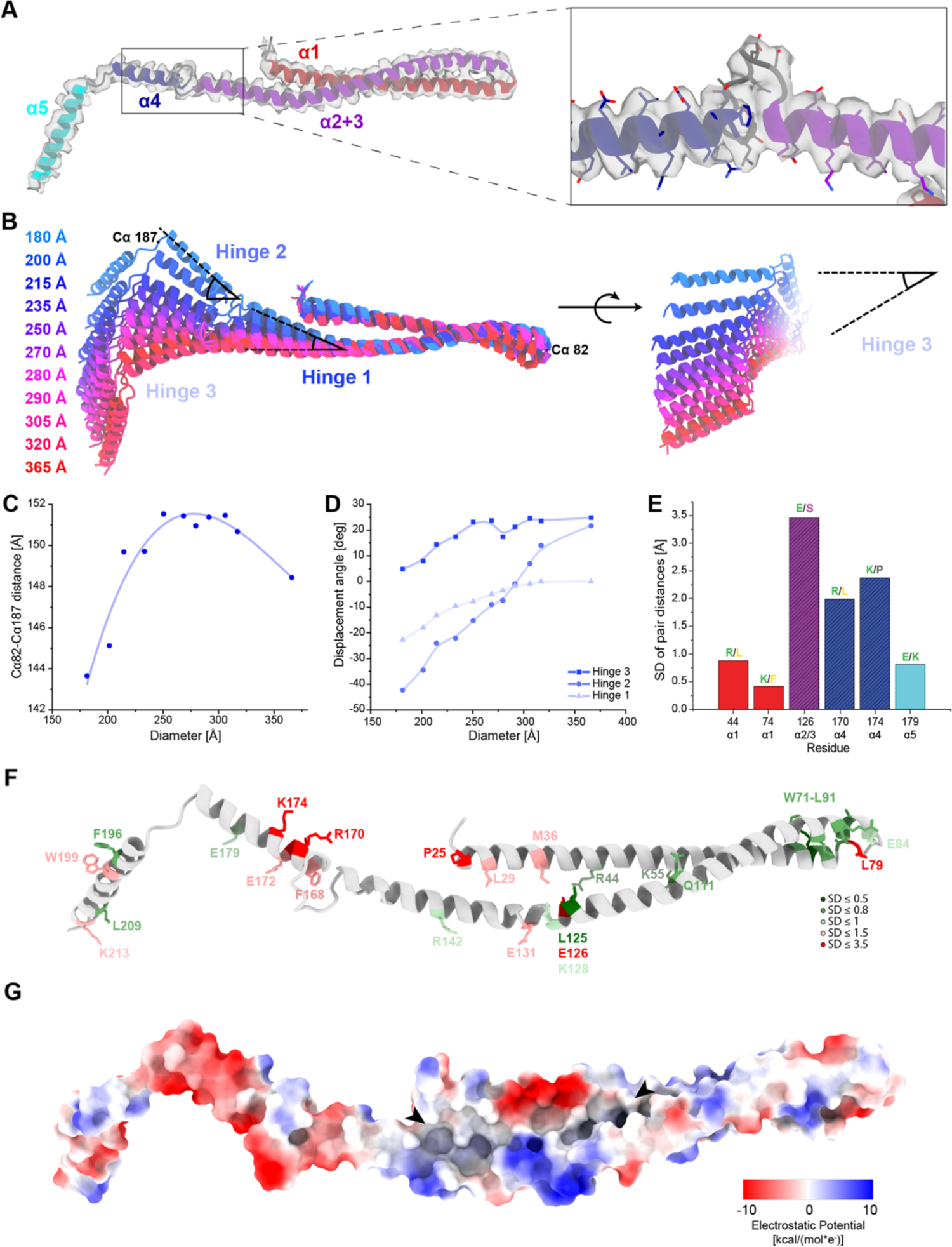
Three hinge regions enable the conformational plasticity of PspA. **A:** Model of a PspA+ATP monomer showing details of the density at the loop connecting α3 and α4 (α1 red, α2+3 violet, α4 blue, α5 cyan) **B:** Superimposed PspA+ATP monomer structures aligned on the hairpin between α1 and α2+3 and color-coded by increasing rod diameter (blue to red for increasing diameters). Hinges 1-3 and the corresponding displacement angle are indicated by opening angle symbols (black). **C:** Scatter plot of PspA+ATP rod diameters over distance between Cα 82 and Cα 187. **D:** Scatter plot of PspA+ATP rod diameter against the displacement angle at Hinges 1-3. **E:** Bar plot of selected pair distances evaluated over all determined diameter assemblies. For a plot of all evolutionarily conserved residue pairs and a more detailed explanation see Suppl. Fig. 3C. Color code for residue letters: green=charged; yellow=hydrophobic; pink=polar+uncharged: grey=special cases. Color code for boxes: α1 red, α2+3 violet, α4 blue, α5 cyan. **F**: Model of PspA+ATP monomer with evolutionarily conserved amino acid residues highlighted and colored according to the flexibility of their contacts (green: low flexibility (SD<1); red: high flexibility (SD>1)). **G**: Model of a PspA+ATP monomer with electrostatic surface coloring. Arrowheads show the start and end of the non-conserved hydrophobic groove.

To examine the contacts between protomers within rod assemblies of different diameters, we next determined the Cα-distance changes over the different diameters between evolutionarily conserved residues of neighboring PspA monomers with respect to the smallest 180 Å diameter and computed the standard deviation as a measure of change (Fig. 3E**, Suppl.** Fig. 3B**/C**). The observed standard deviations fell into two major groups: contacts with distance changes smaller than 1 Å and larger than 1 Å that we assumed to keep fixed contacts or switch contacts with increasing diameters, respectively. The fixed intermolecular contacts consisted of electrostatic as well as hydrophobic interactions. The corresponding residues are clustered in two regions at the hairpin α1/α2-α5 interface (W71, K74, R88, L91, F196, and M212) and the α2+3-α1/α2 hairpin interface (R44, K55, L125, K128, L138 and R142). Additional fixed residues can be found at the start and end of α4 (F168 and E179) (Fig. 3F). Fixed contacts likely serve as anchor points critical for keeping the polymer assembled during structural rearrangements resulting in diameter transitions. In addition to these fixed contacts, we identified several evolutionarily conserved contacts that have large Cα-distance changes and likely switch their interacting residues when comparing different diameters (E126, R170, and K174). Together with a hydrophobic groove in the hairpin between α1 and α2 formed by evolutionarily mostly non-conserved residues (Fig. 3G), they provide the necessary flexibility to allow subunits to slide relative to each other accompanied by the described conformational change of the monomers. Furthermore, the helical rise of the rods decreases with increasing diameters, resulting in a linear mass-per-length increase (Suppl. Fig. 3D-F). In conclusion, the structural plasticity of PspA rods is accomplished by a combination of evolutionarily conserved fixed and switching residue interactions and of non-conserved hydrophobic groove interactions between neighboring PspA molecules.

### PspA rods bind and hydrolyze ATP

Based on the observed modulated structural plasticity in the presence of ATP and the recently identified NTPase activity of the closely related Vipp1 protein (Gupta et al., 2021; Junglas et al., 2020b; Ohnishi et al., 2018; Siebenaller et al., 2021), we investigated whether PspA has a similar ATPase activity. In the resolved PspA cryo-EM structures, we could not identify a canonical P-loop or Walker A motif but conserved residues R44, E126, and E179 whose mutations significantly decreased the ATPase activity of Vipp1 (Gupta et al., 2021) (Fig. 4A). Therefore, we performed a malachite-green based ATPase assay and measured phosphate release in presence of ATP (Fig. 4B) using electrophoretically pure PspA (Suppl. Fig. 4A). The measured ATPase activity of PspA was approx. 10 times lower when compared with *bona fide* AAA+ ATPases, such as *E. coli* PspF (Suppl. Fig. 4B). The ATPase activity was not significantly reduced in absence of Mg^2+^ or even in presence of EDTA, but could be slightly decreased by high concentrations of classical ATPase inhibitors such as ATPγS or AMPPCP (Lacabanne et al., 2020). Interestingly, measuring the activity upon mutating residues R44, E126, and E179 revealed that these residues are critical for the PspA ATPase activity, as found for Vipp1. Single mutations already decreased the ATPase activity by 60-70% and double residue mutations led to a decrease by ∼80%. Simultaneous mutation of all three residues reduced the activity by ∼90%. In contrast, PspA α1-5, *i.e.*, truncated PspA with helix α0 and the first 22 aa removed that are far away from the putative active site, did not significantly reduce the ATPase activity. To verify ADP release after ATP hydrolysis, we performed an ADP-Glo assay and measured an ATPase activity of 3 h^-1^, as determined with the phosphate release assay (Fig. 4C). After incubation with lipid membranes, the ATPase activity was increased by ∼210 %. To further investigate the mode of ATP hydrolysis by PspA, we used mass spectrometry to verify that PspA rods can bind ATP and hydrolyze ATP to ADP. Indeed, after incubation of a PspA with ATP and extraction of the protein bound nucleotides, we identified two peaks by HPLC/MS-MS with *m/z* ratios corresponding to ADP and ATP (Fig. 4D). Interestingly, traces of ADP and ATP have also been found in a PspA sample without addition of ATP, indicating that minor amounts of ADP/ATP were routinely co-purified from the expression host (Suppl. Fig. 4C). In agreement with our phosphate release measurements, HPLC/MS-MS indicated that the R44K/E126Q/E179Q mutant was still capable of ATP binding and hydrolysis, whereas only to about half of the wild-type protein (Suppl. Fig. 4D-E). Notably, the tested mutants were still capable of forming large rod structures comparable with the wild-type protein (Fig. 4E). Biochemical and mass spectrometry analysis of PspA confirmed the conservation of ATPase activity in bacterial PspA and Vipp1 proteins. Together, we showed that our PspA preparations were capable of binding and hydrolyzing ATP albeit with a one order of magnitude lower activity than observed for canonical P-loop or Walker A motif-ATPases.

**Figure 4:**
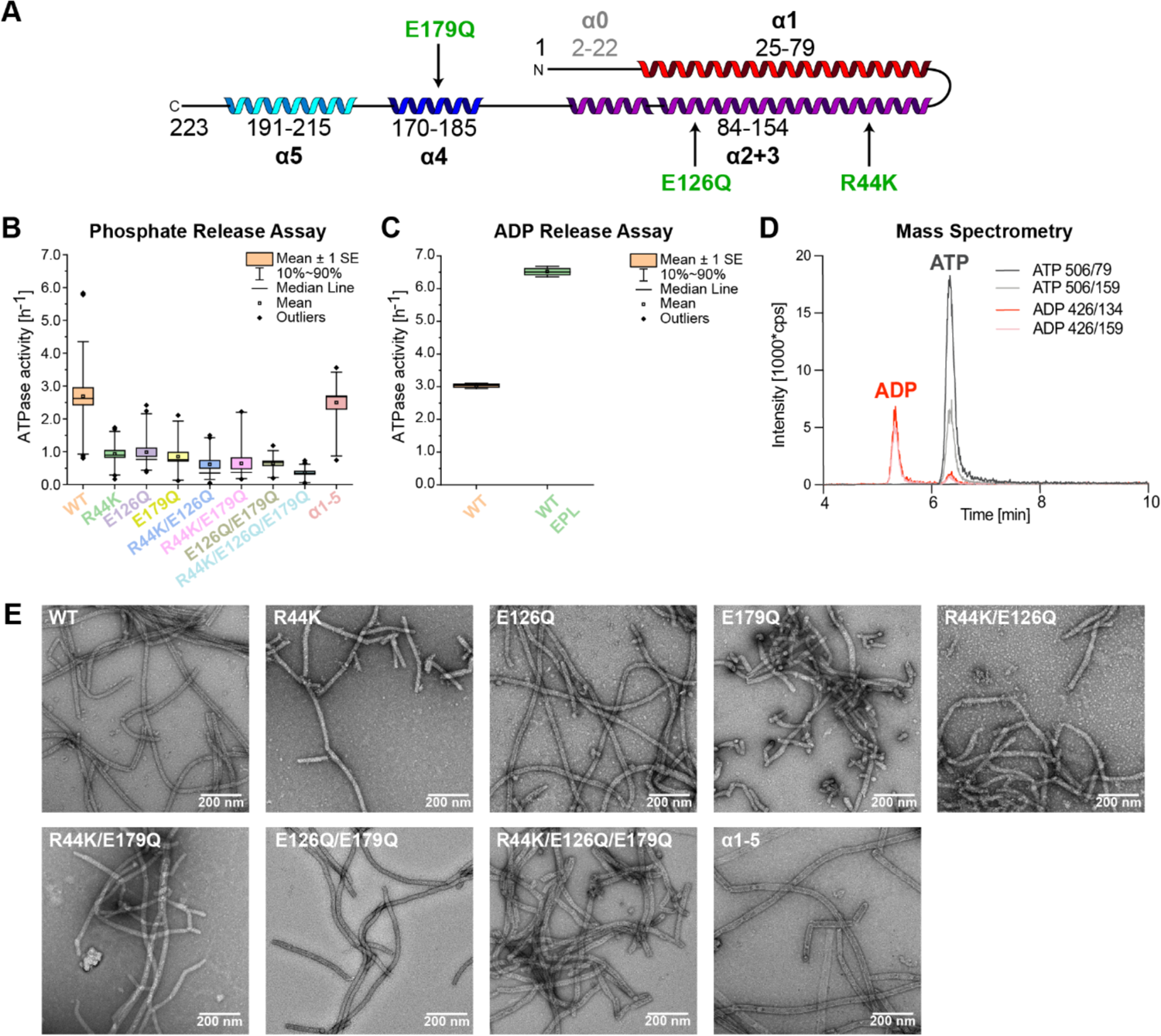
Nucleotide binding and hydrolysis by PspA. **A:** Schematic secondary structure topology of the PspA monomer (α1 red, α2+3 violet, α4 blue, α5 cyan). Relevant mutations are indicated in green. **B:** ATPase activity of PspA wild type (WT) and mutants measured by a malachite-green based assay. The boxplots show the mean ± standard error as boxes, the 10% to 90% percentile as whiskers, and outliers as diamonds. *p-* values of two-sample t-test: *p(WT_vs R44K/E126Q/E179Q)*=7.18*10^-9^, *p(WT_vs α1-5)*=0.63 (n(WT)=36, n(R44K)=24, n(E126Q)=27, n(E179Q)=18, n(R44K/E126Q)=21, n(R44K/E179Q)=18, n(E126Q/E179Q)=18, n(R44K/E126Q/E179Q)=21, n(α1-5)=18; For all measurements, samples of at least 3 biological replicates were included. **C:** ATPase activity of the WT protein and the α1-5 mutant without and in the presence of EPL; WT (orange): ATPase activity without EPL; WT EPL (green): ATPase activity in the presence of EPL (SUVs made from E. coli polar lipid extract); α1-5 variant (purple); n=3. **D:** HPLC/MS-MS of PspA + ATP after 24 h incubation and extensive washing. Different color lines represent different MRM transitions. For ADP, MRM transitions are 426/134 (red) and 426/159 (light red). For ATP MRM transitions are 506/79 (dark grey) and 506/159 (light grey). The ADP fragments below the ATP peak are formed by in-source fragmentation of ATP in the ESI source. **E:** Negative staining EM micrographs of PspA WT protein and PspA mutants. Magnification 57 kx.

### ATP enhances SynPspA-induced membrane remodeling

To assess the consequences of PspA’s structural plasticity in the context of a membrane remodeling activity, we analyzed the cryo-images for structural and morphological changes of the added EPL SUVs and the effect of ATP under reconstitution conditions developed for optimal tubulation (50 nm SUVs and *in situ* refolding) (Fig. 5A **and B**). In the absence of PspA, control SUVs had a narrow bilayer thickness distribution from 27 to 32 Å with a mean bilayer thickness of 31 Å (Fig. 5C). After incubation with PspA, we observed PspA rods engulfing and tubulating membrane. Notably, PspA rods engulfing membranes did not have uniform diameters. Instead, they were frequently attached to vesicles with one wider end tapering towards the distal end (Fig. 5B). Moreover, engulfed membranes were continuous lipid tubes as well as small, isolated vesicles and membrane patches. After incubation with PspA, SUV membranes were significantly thicker on average, with a broad distribution and a major peak at 38 Å and a minor peak at 68 Å. In the presence of PspA and ATP, the bilayer thickness distribution closely resembles the PspA sample (mean thickness 37 Å). The observed increase in bilayer thickness after incubation with PspA is consistent with a previous report (Junglas et al., 2021). To monitor changes in vesicle sizes, we used the vesicle perimeter as a descriptor (Fig. 5D**, Suppl.** Fig. 5A). The EPL SUVs alone had a mean perimeter of 100 nm corresponding to a mean diameter of 32 nm. Similar to the bilayer thickness, the distribution of vesicle perimeters was significantly increased in the presence of PspA and PspA+ATP, peaking at 160 nm and 150 nm, respectively. The vesicle size increase observed after incubation with PspA agrees with the previously reported PspA-mediated vesicle fusion (Junglas et al., 2021). Another prominent feature of the vesicles containing PspA was the formation of small, double-membrane vesicles (*i.e*., small vesicles that are encapsulated by a larger vesicle), in analogous topology to intra-luminal vesicles (ILVs) (Fig. 5B, red arrowheads). Approx. 9 % of all vesicles were double-membrane vesicles in the control whereas the share of double-membrane vesicles was increased in the PspA preparations to 41 % and was highest in the PspA+ATP sample with 52 % (Fig. 5E). Using the ATPase-deficient mutant PspA R44K/E126Q/E179Q as a negative control, we validated that indeed ATPase activity of PspA and not the addition of ATP gave rise to the increased number of double-vesicles in the PspA+ATP sample. In addition, the distance between the two enclosed vesicle membranes, termed here the enclosure distance, was on average higher in the control vesicles (>70 Å) with a very broad enclosure distance distribution (Suppl. Fig. 5A-C). Both, the PspA as well as the PspA+ATP sample, showed a well-defined distance distribution with a mean enclosure distance of 67 and 62 Å, respectively. Compared with the vesicle perimeter, only the smallest vesicles (perimeter 100 – 200 nm) showed this constant enclosure distance of ∼50 – 70 Å (Suppl. Fig. 5D). Together, quantitative image analysis of vesicle characteristics showed that the presence of ATP increases the occurrence of double-membrane vesicles by enhanced membrane remodeling capabilities of PspA.

**Figure 5:**
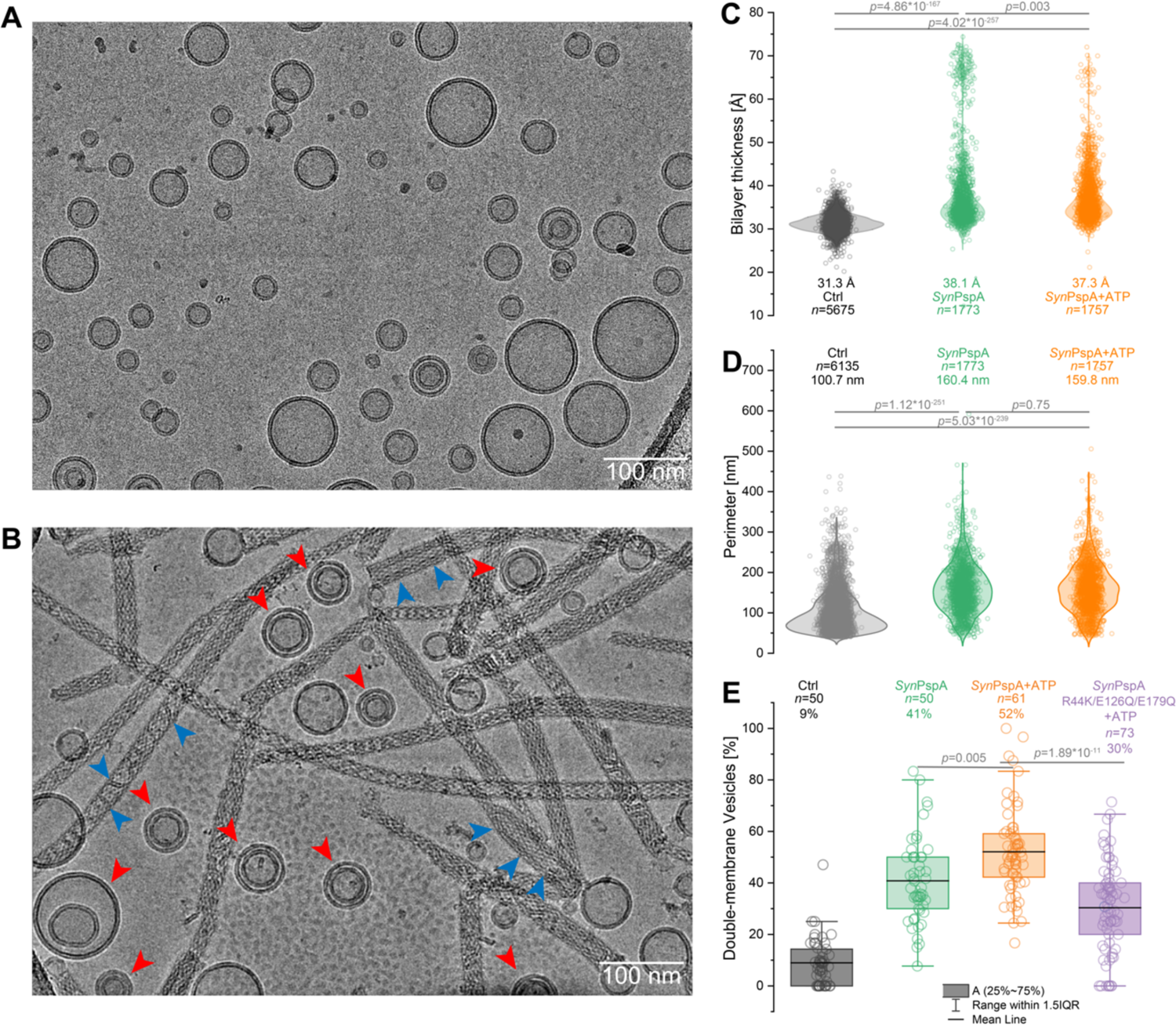
Changes of EPL vesicle morphologies after *Syn*PspA ± ATP reconstitution. **A:** Example cryo-EM micrograph of EPL control vesicles. **B:** Example cryo-EM micrograph of the PspA+EPL sample. Red arrowheads indicate encapsulated vesicles. Blue arrowheads indicate engulfed membranes. **C:** Violin plot of the bilayer thickness of control EPL SUVs (grey), PspA+EPL SUVs (green), and PspA+EPL+ATP SUVs (orange). The mean bilayer thickness, the number of measured vesicles *n*, and *p* values from a two-sample t-test are indicated on the graph. **D:** Violine plot of the vesicle perimeter of control EPL SUVs (grey), PspA + EPL SUVs (green), and PspA+EPL +ATP SUVs (orange). The mean vesicle perimeter, the number of measured vesicles *n*, and *p* values from a two-sample t-test are indicated on the graph. **E:** Box plot of the relative occurrence of double-membrane vesicles in each sample: control EPL SUVs (grey), PspA+EPL SUVs (green), PspA+EPL+ATP SUVs (orange), and PspA(R44K, E126Q, E179Q)+EPL+ATP SUVs (violet). The relative number of double-membrane vesicles, the number of measured vesicles *n*, and *p* values from a two-sample t-test are indicated on the graph (Ctrl *vs. Syn*PspA: p=1.83*10^-19^, Ctrl *vs. Syn*PspA + ATP: p=1.13*10^-31^, Ctrl *vs. Syn*PspA R44K/E126Q/E179Q + ATP: p=1.24*10^-14^, *Syn*PspA *vs. Syn*PspA + ATP: p=0.005, *Syn*PspA + ATP *vs. Syn*PspA R44K/E126Q/E179Q + ATP: p=1.89*10^-11^).

### Effect of non-hydrolyzable ATP analogs and mutations on PspA’s structural plasticity

To further investigate how nucleotide binding and hydrolysis affects the PspA rod diameter distribution, we next analyzed the rod diameter distribution after addition of ATP to preformed PspA rods using cryo-EM. The resulting rod distribution showed two distinct peaks at 215 Å and 250 Å diameters whereas only a minor shift to larger diameters, compared with the rod distribution after the addition of ATP during rod assembly, was observed (Suppl. Fig. 6A **and B**). These observations indicate that only minor diameter changes are possible once the rods have formed and larger changes can only occur during rod assembly or re-assembly. Additionally, we tested the effect of the non-hydrolyzable ATP analogs AMPPCP and ATPγS on the diameter distribution using cryo-EM. First, upon addition of AMPPCP during rod formation, we found that the diameter distribution is approximately indistinguishable from the distribution in the presence of ADP supported by the average ± standard deviation measurements: 240±40 Å and 240±50 Å of the ADP and the AMPPCP distribution, respectively (Suppl. Fig. 6C **and D**). Unlike for canonical ATPases (Krasteva and Barth, 2007), these data indicate that AMPPCP mimics the ADP-bound state in the case of PspA. Second, upon addition of ATPγS, the diameter distribution overlapped highly with the ATP sample although that showed an additional tail of higher diameter rods (>300 Å) (Suppl. Fig. 6E **and F**). Assuming that ATPγS mimics the ATP transition state, this observation suggests that complete ATP turnover rather than mere binding is required for the formation of rods with large diameters.

In the series of determined PspA structures we found the second hinge, *i.e.,* the loop connecting helices α3 and α4 (loop α3/α4), to be critically important for the conformational adjustments of PspA monomers in rods with higher diameters and also participating in the putative nucleotide binding region. Therefore, we generated a PspA mutant lacking ten residues of the loop E156 – S165 (PspA dL10) (Fig. 6A) and analyzed the ATPase activity as well as diameter distribution. First, PspA dL10 showed an ATPase activity reduced by over 90% compared with wild-type PspA, although the above mutated residues critical for the ATPase activity (R44, E126, E179) were still intact (Fig. 6B). Second, we employed cryo-EM to determine the structure of PspA dL10 and compared it with the wild-type protein. We found that PspA dL10 still formed rods with symmetries and shapes also present in the wild-type sample, although with a narrower distribution centered at slightly higher diameters (Fig. 6C **and D**). Upon addition of ATP during the formation of PspA dL10 rods, we did not observe significant differences in the diameter distribution (220±20 Å (dL10) *vs*. 230±30 Å (dL10 ATP), in agreement with the highly reduced ATPase activity (Fig. 6C **and E****, Suppl. Fig. G and H**). The detailed cryo-EM structures of PspA dL10 rods were almost identical to the wild-type rods, while the density for the loop between helices α3 and α4 was missing (Fig. 6F). Together, the dL10 rod structures indicate that loop α3/α4 is not essential for the formation of PspA rods *per se*, but critical for the formation of rods with very low and high diameters, as well as for the associated ATPase activity. By removing the loop α3/α4, we reduced the conformational freedom providing an efficient means to restrict the observed structural plasticity of PspA assemblies.

**Figure 6:**
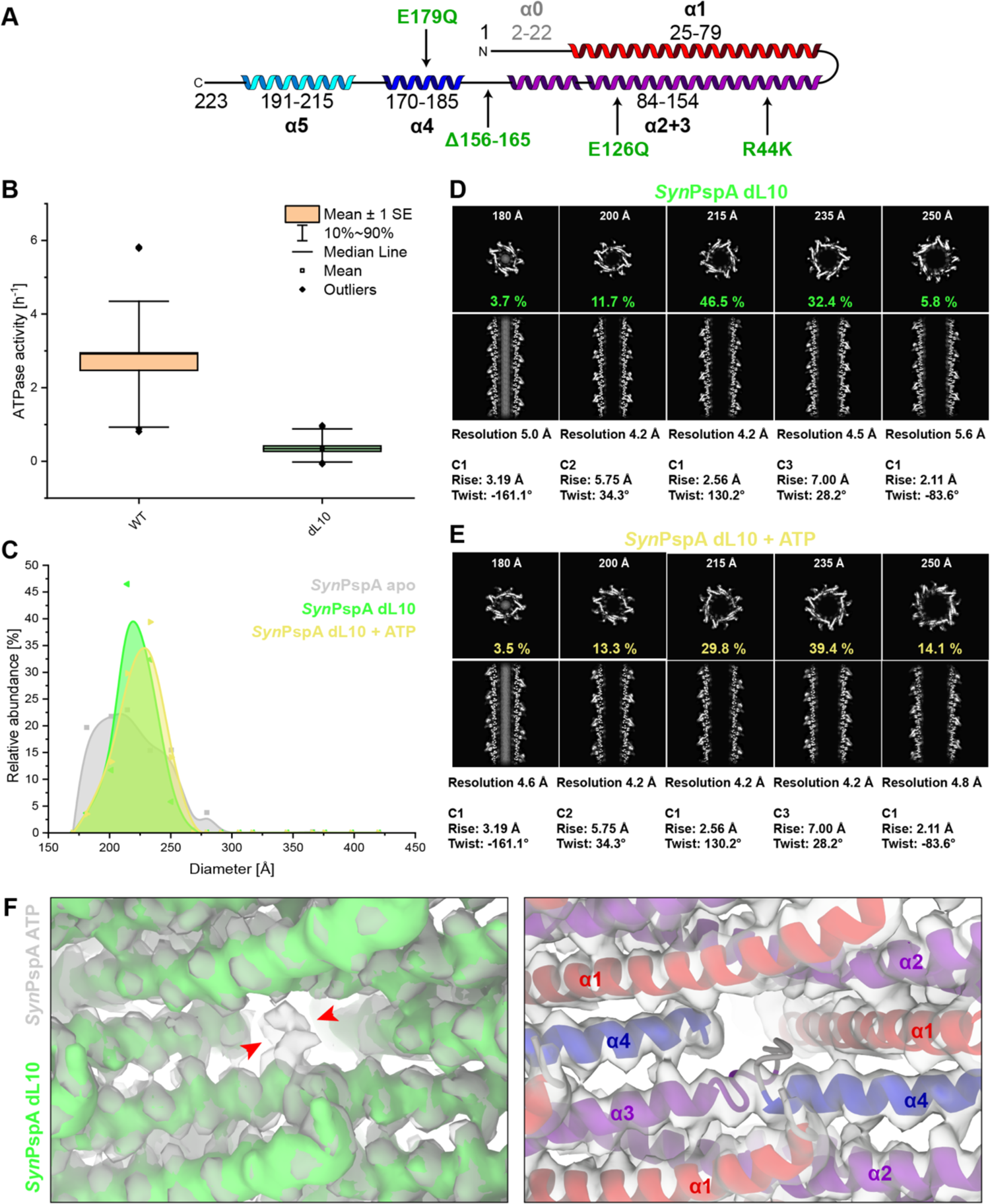
ATPase activity and structure of the PspA dL10 mutant. **A:** Schematic secondary structure topology of the PspA monomer (α1 red, α2+3 violet, α4 blue, α5 cyan). The deletion of loop residues 156-165 (dL10) and relevant mutations are indicated in green. **B:** ATPase activity of PspA wild type (WT) and dL10 measured by a malachite-green based assay. The boxplots show the mean ± standard error as boxes, the 10% to 90% percentile as whiskers, and outliers as diamonds. *p-*Values of two-sample t-test: *p(WT_vs dL10)*=3.92*10^-10^, (n(WT)=39, n(dL10)=23; For all measurements, samples of at least 3 biological replicates were included) **C:** Histogram of rod diameters and the normalized abundance of PspA dL10 rods (green), PspA dL10 rods in the presence of ATP (yellow), and PspA WT rods (grey) for comparison. PspA dL10 has a narrow diameter distribution that is not affected by ATP addition. **D, E:** The associated cryo-EM structures of PspA dL10 and PspA dL10 + ATP are shown in cross-sectional top and side views, respectively. **F: Left:** Comparison of the cryo-EM density maps of PspA+ATP (grey) and PspA dL10 (green). Red arrowheads indicate the additional density for the loop that is missing in the dL10 map. **Right:** Cryo-EM density map of PspA dL10 with the PspA WT model.

## Discussion

In this work, we revealed in molecular detail the structural plasticity of PspA that is capable of forming a large series of related assembly structures. We determined a total of 61 cryo-EM structures of helically assembled PspA rods spanning rod diameters from 180 to 400 Å under different conditions, *i.e.,* in the absence of nucleotides, or the presence of ADP and ATP, respectively (**Figs. 1, 2**). We found that the distribution of PspA rod diameters is affected by the presence of the nucleotides, resulting in the largest rod diameters and widest diameter range after incubation with ATP, intermediate-sized diameter rods with narrow distribution in the presence of ADP, and, in the absence of any nucleotide, the small diameter rods with intermediate distribution were observed. In total, we obtained 14 unique assembly structures over the five samples including non-hydrolyzable ATP analogs AMPPCP and ATPγS. The changes of the PspA polymer diameters were caused by conformational changes primarily at three hinge regions between the α-helical segments of PspA (Fig. 3). Moreover, we biochemically characterized the ATPase activity of PspA and studied mutants abrogating the hydrolytic activity while maintaining rod formation (Fig. 4). To link the structural plasticity and ATPase activity of PspA with a potential function, we analyzed changes in the membrane vesicle morphology induced by PspA in the absence and presence of ATP and found that ATP appears to enhance the membrane remodeling capabilities of PspA (Fig. 5). Structural and biochemical analysis of a mutant lacking the loop connecting helix α3 and α4 revealed that this hinge region is essential for the structural plasticity and ATPase activity of PspA (Fig. 6).

Superimposing the now-determined structures, we identified three hinge regions that enable the PspA conformation to flex from a high-tension bow shape over a low-tension linear shape back to a high-tension inverted bow shape. These molecular shape changes expose slightly modified subunit interfaces and, as a result, PspA mass packing per unit length of polymer increases. Ultimately, these packing changes give rise to increasing polymer diameters, from the observed lowest diameters of 180 Å to the highest observed diameters up to 400 Å. As higher diameter rods only formed in the presence of ADP or ATP, the monomer tension in these rods may be stabilized in the process of ATP binding/hydrolysis, presumably due to the formation of new interactions between protein and the nucleotide. Interestingly, the putative ATP/ADP binding pocket is formed by four PspA chains (j_0_, j_-1_, j_+26_, and j_+27_) and α1-4 as well as the respective loop regions. Relevant catalytic residues, including residues of helices α3 and α4 connecting loop point toward the potential binding site and were found to be solvent accessible. The three hinge regions are close to the putative nucleotide binding residues in the polymeric assembly. According to the presence of weak density in a few or the lack of density in most cryo-EM reconstructions and in reference to the Vipp1 structure (Gupta et al., 2021), it is likely that not every binding site is equally occupied by nucleotides. In the determined Vipp1 ring structures of cyclical symmetry, ADP was found in one discrete Vipp1 layer, *i.e.*, the most constricted layer of the ring. The presence of ADP in this single layer suggests that only a fraction of putative sites correspond to be active in one ring assembly. The low ATPase activity of PspA from *Synechocystis* is in accordance with the reported GTPase/ATPase activity of the closely related Vipp1 and PspA from *E.coli* (Gupta et al., 2021; Junglas et al., 2020b; Ohnishi et al., 2018; Siebenaller et al., 2021). In support, low competition rates with well-characterized non-hydrolyzing ATP analogs reveal differences in ATP turnover when compared with canonical P-loop or Walker A motif-ATPases. However, when only few layers per rod molecule are considered actively hydrolyzing, the effective turnover can be much higher locally. For example, considering the average length of PspA rods are 2.6 ± 1.3 µm, a total of approx. 10,000 monomers are found in rod corresponding to 1000 turns depending on the symmetry. Therefore, we reason that local ATP turnover can be approx. 1000 times higher, *i.e.*, three orders of magnitude, resulting apparent 50 ATP molecules per min (3000 h^-1^), which is in the determined activity range of actively remodeling dynamin (Doo Song et al., 2004).

Interestingly, ATP hydrolysis is critical for the disassembly or assembly of eukaryotic ESCRT-III polymers mediated by the dedicated AAA+ ATPase Vps4 (Mierzwa et al., 2017; Nickerson et al., 2006). By hydrolyzing ATP, Vps4 can remove and unfold individual ESCRT-III subunits from the polymer structures (Yang et al., 2015). Although some Psp systems also have an AAA+ ATPase (PspF), it does not appear to be essential for the membrane remodeling properties of PspA, as not all organisms that contain PspA also contain PspF (*e.g., Synechocystis* sp. PCC 6803 does not). Vps4 is thought to recruit ESCRT-III members via its N-terminal microtubule interacting and trafficking (MIT) domain (Obita et al., 2007; Stuchell-Brereton et al., 2007). Vps4 triggers a conformational change in ESCRT-III polymers by exchanging subunits of the filament polymers to allow greater curvature, which can cause the tubes to constrict (Caillat et al., 2019). This conformational change can lead to the fission of bound vesicles by decreasing filament diameter in membrane-bound complexes (Chiaruttini and Roux, 2017). For initial ILV biogenesis, the complete ESCRT-III heteropolymer structure is assembled, and after binding of Vps4, membrane constriction and release of the ILVs can occur (Adell et al., 2014). However, it remains to be established whether the observed modulation of PspA rod diameters through ATP/ADP binding and/or hydrolysis is of physiological relevance in the bacterial cell. PspA rods mixed with liposomes produced double-membrane vesicles that share the same topology as ILVs and are likely a result of inward-vesicle budding, *i.e.,* an intraluminal budding process (Junglas et al., 2021). In accordance with our previous work, we suggest that double-membrane vesicles are formed by cross-membrane vesicle transfer, *i.e.,* by releasing of PspA-engulfed membrane into an acceptor vesicle. In the current work, we demonstrated that after PspA incubation the number of double-membrane vesicles increased in the presence of ATP. This observation indicates that the membrane remodeling activity of PspA is stimulated by ATP, *i.e.,* by facilitating the uptake of vesicle membranes when larger diameter rods engulf membranes more efficiently than low diameter rods due to lowered membrane curvature. Given these observations, we can now directly link the elusive NTPase activity of the Vipp1/PspA protein family (Gupta et al., 2021; Junglas et al., 2020b; Ohnishi et al., 2022, 2018; Siebenaller et al., 2021) to an enhanced production of ILV-like vesicles.

In the case of eukaryotic ESCRT-III proteins, Vps4 has been proposed to mediate constriction and membrane fission accompanied by a reduction in the diameter of the polymeric structures through the hydrolysis of ATP (Maity et al., 2019; Schöneberg et al., 2018), in analogy to dynamin that accomplishes constriction through the hydrolysis of GTP. In contrast, ATP addition during PspA tube formation leads to dilation of rod diameters. The results of ATP incubation with preformed PspA rods suggest that large diameter changes of PspA rods can apparently only occur during rod assembly or re-assembly, which may be accomplished by constant disassembly and assembly by chaperones or PspF in the bacterial cell, in analogy to Vps4 in eukaryotic cells. While it cannot be clearly deduced from our data whether ATP binding and/or hydrolysis leads to the dilation (or ADP/ATP release leads to rod constriction), the structural similarity between PspA and the ESCRT-III proteins raises the question of whether the ESCRT-III proteins themselves also possess a nucleotide binding pocket similar to that of PspA and therefore have an intrinsic ATPase activity. Reports are emerging that eukaryotic as well as archaeal ESCRT-III proteins can bind DNA or have been localized in foci close to chromatin (Nachmias et al., 2022; Talledge et al., 2018; Vietri et al., 2020). A cryo-EM structure of an IST1/CHIMP1B polymer bound to negatively charged DNA, deposited on a preprint server, revealed a binding site at the inner lumen of the tubes in potential competition with the negatively charged lipid membrane (Talledge et al., 2018). In contrast, the here-identified putative ATP binding pocket in PspA polymers is located centrally in the tube wall, sandwiched between adjacent subunits. Together, a common feature of bacterial and eukaryotic ESCRT-III proteins (together with associated AAA+-ATPases such as Vps4) is the ability of binding nucleotides and the conformational changes of the monomers.

In this study, we describe how a combination of fixed and switching residue interactions potentially mediate the observed structural plasticity of PspA. The fixed residues are of polar as well as hydrophobic nature and likely serve as anchor points to keep the polymer assembled. Switching residues include evolutionarily non-conserved hydrophobic residues forming a groove on the helix α1/α2 hairpin, which are responsible for establishing the required structural flexibility of assembling different diameter rods. Defined residue contacts also explain why discrete diameter states were observed as opposed to an ensemble of quasi-continuous changes of diameters. The here observed structural plasticity is a common property of several polymeric protein assemblies in various biological fields. For instance, structural rearrangements were already identified in early fundamental work on tobacco mosaic virus (TMV) coat protein when pH and ionic strength determined the type of disc, stacked discs, and helix assemblies (Klug, 1999). Other viruses, such as mumps nucleocapsid viruses, also exhibit such a structural plasticity as they were observed in a series of different assemblies varying their number of helical units resulting in denser and looser packing of the viral genome (Shan et al., 2021). Smaller helical units from short synthetic peptides are commonly found in different packing arrangements (Egelman et al., 2015). Moreover, one of the structural hallmarks of neurodegenerative diseases is a structural plasticity in the form of varying folds and different protofilament packings within a common cross-β structure framework of amyloid fibrils (Shi et al., 2021). Structural plasticity has also been observed in a series of lipid remodeling complexes. For instance, ANTH/ENTH clathrin adaptor complexes have been shown to form helical assemblies over a range of different diameters each varying by an additional subunit in the helical turn (Skruzny et al., 2015). Moreover, eukaryotic ESCRT-III proteins have been shown to portray a structural polymorphism assembling in sheets, strings, rings, filaments, tubules, domes, coils, and spirals (Huber et al., 2020; McCullough et al., 2015; Pfitzner et al., 2021, 2020). In fact, the structural changes observed now for the hinges connecting helices α3 and α4 as well as helices α2 and α3 are consistent with structures of 180 Å CHMP1B/IST1 copolymer and 280 Å diameter CHMP1B filaments showing IST1 binding-induced conformational changes in the homologous hinge regions of the ESCRT-III protein CHMP1B (Nguyen et al., 2020). Similarly, tubular assemblies with varying diameters were observed with human ESCRT-III CHMP2A/CHMP3 assemblies, and it has been hypothesized that such plasticity is involved in filament sliding that could drive membrane constriction (Azad et al., 2023). A related membrane remodeling molecule, *i.e.*, human dynamin-1, has been shown to exhibit large-scale GTPase-mediated structural rearrangements accompanied by a change in assembly diameter (Liu et al., 2021a). In the case of the canonical dynamin, GTP binds to the dedicated catalytic so-called G-domain and enables filament constriction for the final membrane fission step. In contrast, PspA binds and hydrolyzes ATPs between the subunits of the polymer presumably causing dilation of rods. However for EHD1, a dynamin family member, it has been shown that ATP hydrolysis mediates membrane remodeling by dilation of membrane tubulation EHD1 scaffolds (Deo et al., 2018). Although the here observed PspA’s structural plasticity may be crucial for the PspA membrane remodeling activity (Junglas et al., 2021), it remains to be seen whether the small-scale conformational changes are sufficient to trigger membrane dilation/constriction. Clearly, further characterization is required, in particular, to better understand the structural transitions of PspA assemblies in the presence of lipid membranes.

## Material and Methods

### Expression and purification of PspA and EcoPspF

PspA wild-type (wt) (*slr1188*) of *Synechocystis* sp. PCC 6803 and associated mutants (R44K, E126Q, E179Q, R44K/E126Q, R44K/E179Q, E126Q/E179Q, R44K/E126Q/E179Q, α1-5 (deletion of α0 (aa 1-23), dL10 (deletion of aa 156-165, the loop between α3 and α4)) were heterologously expressed in *E. coli* C41 cells in TB medium using a pET50(b) derived plasmid.

For purification of PspA and associated mutants under denaturing conditions, cells were resuspended in lysis buffer containing 6 M urea (10 mM Tris-HCl pH 8.0, 300 mM NaCl) supplemented with a protease inhibitor. Cells were lysed in a cell disruptor (TS Constant Cell disruption systems 1.1 KW; Constant Systems, Daventry, UK). The crude lysate was supplemented with 0.1% (v/v) Triton X-100 and incubated for 30 min at RT. Subsequently, the lysate was cleared by centrifugation for 15 min at 40,000 g. The supernatant was applied to Ni-NTA agarose beads. The Ni-NTA matrix was washed with lysis buffer and lysis buffer with additional 10 – 20 mM imidazole. The protein was eluted from the Ni-NTA with elution buffer (10 mM Tris-HCl pH 8.0, 1000 mM imidazole, 6 M urea). The fractions containing protein were pooled, concentrated (Amicon Ultra-15 centrifugal filter 10 kDa MWCO), and dialyzed overnight against 10 mM Tris-HCl pH 8.0 or buffer containing 2 mM ADP or ATP with 4 mM MgCl_2_ (8 °C, 10 kDa MWCO) including three buffer exchanges. Protein concentrations were determined by measuring the absorbance at 280 nm of PspA diluted in 4 M guanidine hydrochloride using the respective molar extinction coefficient calculated by ProtParam (Gasteiger et al., 2005).

PspF(1-275) of *E. coli* K12 was heterologously expressed in *E. coli* BL21 (DE3) cells in TB medium using a pET50(b) derived plasmid. For purification of EcoPspF, cells were resuspended in lysis buffer (10 mM Tris-HCl pH 8.0, 5% (v/v) Glycerol, 50 mM NaCl) supplemented with a protease inhibitor. Cells were lysed in a cell disruptor (TS Constant Cell disruption systems 1.1 KW; Constant Systems) followed by centrifugation of the lysate for 15 min at 5,000 g. The supernatant was applied to Ni-NTA agarose beads. The Ni-NTA matrix was washed with lysis buffer and lysis buffer with 20 - 60 mM imidazole. The protein was eluted from the Ni-NTA with elution buffer (10 mM Tris-HCl pH 8.0, 50 mM NaCl, 1,000 mM imidazole). The fractions containing protein were pooled and dialyzed overnight against dialysis buffer (10 mM Tris-HCl pH 8.0, 5% (v/v) Glycerol, 50 mM NaCl, 0.1 mM EDTA, 1mM DTT) (8 °C, 10 kDa MWCO) including three buffer exchanges. Protein concentrations were determined by measuring the absorbance at 280 nm of EcoPspA using the respective molar extinction coefficient calculated by ProtParam (Gasteiger et al., 2005).

### Liposome preparation and lipid handling

Chloroform dissolved *E. coli* polar lipid (EPL) extract was purchased from Avanti polar lipids. Lipid films were produced by evaporating the solvent under a gentle stream of nitrogen and vacuum desiccation overnight. The lipid films were rehydrated in 10 mM Tris-HCl pH 8.0 by shaking for 30 mins at 37 °C. The resulting liposome solution was subjected to five freeze-thaw cycles, combined with sonication at 37 °C in a bath sonicator. SUVs (small unilamellar vesicles) were generated by extrusion of the liposome solution through a porous polycarbonate filter (50 nm pores). For *Syn*PspA lipid reconstitution, unfolded *Syn*PspA (in 6 M Urea), or buffer with 6M Urea for the controls was added to EPL SUVs and incubated at RT for 15 min. Then the mixture was dialyzed overnight against 10 mM Tris-HCl pH 8.0 (8 °C, 10 kDa MWCO) including three buffer exchanges.

### ATPase activity determination

To test phosphate release from PspA upon ATP hydrolysis, a malachite-green assay was used (PiColorLock Gold Phosphate Detection Kit). PspA (4 - 8 µM) was mixed with 2 mM ATP and 4 mM MgCl_2_ in 10 mM Tris-HCl pH 8.0 and incubated at 37 °C for 60 min. The reaction was stopped by the addition of the malachite-green mixture (“Gold mix”) and incubated for 5 min at RT before adding the “stabilizer”. Samples were transferred to a 96-well plate and incubated for 10 min at 25 °C with shaking. Absorbance at 620 nm with pathlength correction was measured using a Tecan infinite M nano plate reader. To determine the P_i_ released by PspA, the absorbance was corrected for ATP and protein background. Then the P_i_ release was calculated by linear regression from a phosphate standard curve. The ATPase activity of PspA was calculated as:

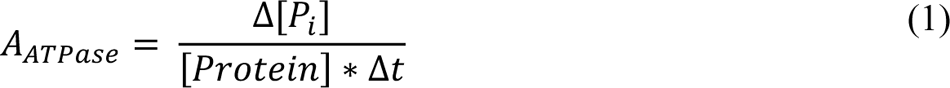

with the average ATPase activity *A_ATPase_*, the phosphate release in µM *Δ[P_i_]*, the protein concentration in µM *[Protein],* and the incubation time of the reaction in hours *t*.

To determine the ATPase activity of PspA in the presence of lipids, the ADP formed during the ATPase reaction of PspA was measured using a Promega ADP Glo Assay. PspA (4 - 8 µM) reconstituted or incubated with lipids was mixed with 1 mM ATP and 2 mM MgCl_2_ in 10 mM Tris-HCl pH 8.0 and incubated at 37 °C for 60 min. After the reaction, “ADP-Glo Reagent” was added and the mixture was incubated for 40 min at 25 °C with shaking to deplete the remaining ATP. Finally, the “Kinase Detection Reagent” was added to detect the released ADP by a Luciferase reaction. After 60 min reaction time at 25 °C, the mixture was transferred to a 96-well plate and luminescence was detected using a BioRad MP Gel-Doc. Luminescence intensity was determined densitometrical using ImageJ (Rueden et al., 2017). ADP concentrations were calculated by linear regression from an ADP standard curve. The ATPase activity was calculated analogously to equation (1).

### Detection and Quantification of ATP/ADP by mass spectrometry

To determine binding of nucleotides to PspA, mass spectrometric analysis was performed. A 2 mL preparation containing 80 nmol PspA, 2 mM ATP and 4 mM MgCl2 in 10 mM Tris-HCl buffer (pH 8.0) was incubated at 37°C (600 rpm) for 24 hours. After incubation, the protein was pelleted at 20,000g (4°C) for 15 minutes. The supernatant was discarded and the protein pellet washed twice with 2 mL of Tris-HCl buffer pH 8.0. After the second wash, the nucleotides were extracted using methanol. The sample was precipitated with 1 mL 80 % methanol with 10 mM Tris-HCl (pH 8.0) for 30 min at room temperature. The extracted nucleotides were separated from the precipitated protein by centrifugation at 20,000 g for 15 minutes (4°C) followed by storage of the extract at −20°C. As a control, a similar nucleotide extraction was carried out with protein directly after purification. To account for autohydrolysis of the nucleotides, 2 mM ATP was incubated for 24 hours at 37°C at the same buffer conditions. The autohydrolysis did not exceed 1%.

The supernatants were dried in a SpeedVac (RVC-2-25, Christ, Osterode, Germany) and resuspended in 100 µl 20 mM triethylammonium acetate pH 9.0. For quantification of ATP/ADP, an HPLC-MS/MS method with ESI in negative modus was developed using a Qtrap6500 instrument (ABSciex, Darmstadt, Germany) coupled with an Agilent 1260 HPLC (Agilent, Waldbronn, Germany). Chromatographic separation was performed on a Zorbax Eclipse Plus (4.6 × 100 mm; 2.6 µm particle size; Agilent, Waldbronn, German). The column temperature was kept at 30°C. The mobile phases consisted of 20 mM triethylammonium acetate pH 9.0 (solvent A) and 20 mM triethylammonium acetate pH 9.0 with 20 % acetonitrile (solvent B) at a flow rate of 500 µL min^-1^. The sample injection volume was 10 µL. An isocratic step of 20 % B (0 to 8 min) was followed by an increase to 99 % B within 0.1 min, which was held for 5 min for cleaning. The gradient returned to 20% B within 0.1 min and the system was equilibrated for 5 minutes. Quantitation after HPLC separation was performed using ESI-MS/MS detection in multiple reaction-monitoring (MRM) mode that allows monitoring a specific fragmentation reaction at a given retention time. The parameters used for the quantification of ATP and ADP are listed in **Suppl. Table 1**. Mass spectrometer settings were: curtain gas (N_2_) 40 arbitrary units (a.u.), the temperature of the source 500 °C, nebulizer gas (N_2_) 80 a.u., heater gas (N_2_) 40 a.u., and ion spray voltage (IS) −4500 V. Data acquisition and processing were carried out using the software Analyst 1.6.1 (ABSciex, Darmstadt, Germany). For quantification, the software Multiquant (ABSciex, Darmstadt, Germany) was used. The calibration curve showed linearity between 10 nM and 10 µM ATP/ADP with a correlation coefficient of R^2^ = 0.9993 (ATP) and R^2^ = 0.9991 (ADP).

### Negative staining electron microscopy

For negative-staining electron microscopy, 3 µL of the sample was applied to glow-discharged (PELCO easiGlow Glow Discharger, Ted Pella Inc.) continuous carbon grids (CF-300 Cu, Electron Microscopy Sciences). The sample was incubated on the grid for 1 min. Then the grid was side-blotted using filter paper, washed with 3 µL water, stained with 3 µL 2% uranyl acetate for 30 s, and air-dried. The grids were imaged with a 120 kV Talos L120C electron microscope (ThermoFisher Scientific/FEI) equipped with a CETA camera at a pixel size of 2.49 Å/pixel (54 kx magnification) at a nominal defocus of 1.0 to 2.5 µm.

### Electron cryo-microscopy

PspA grids were prepared by applying 3.5 μL PspA (**Suppl. Table 2**) to glow-discharged (PELCO easiGlow Glow Discharger, Ted Pella Inc.) Quantifoil grids (R1.2/1.3 Cu 200 mesh, Electron Microscopy Sciences). The grids were plunge-frozen in liquid ethane using a ThermoFisher Scientific Vitrobot Mark IV set to 90% humidity at 10 °C (blotting force −5, blotting time 3 to 3.5 s) or a Leica EM GP2 with sensor assisted back-side (blotting time 3 to 5 s) with the same temperature and humidity settings. Movies were recorded in underfocus on a 200 kV Talos Arctica G2 (ThermoFisher Scientific) electron microscope equipped with a Bioquantum K3 (Gatan) detector operated by EPU (ThermoFisher Scientific).

### Image processing and helical reconstruction

Movie frames were gain-corrected, dose-weighted, and aligned using cryoSPARC Live (Punjani et al., 2017). Initial 2D classes were produced using the auto picker implemented in cryoSPARC Live. The following image processing steps were performed using cryoSPARC. The best-looking classes were used as templates for the filament trace. The resulting filament segments were extracted with 600 px box size (Fourier cropped to 200 px) and subjected to multiple rounds of 2D classification. The remaining segments were reextracted with a box size of 400 px (Fourier cropped to 200 px) and subjected to an additional round of 2D classification. The resulting 2D class averages were used to determine filament diameters and initial symmetry guesses in PyHI (Zhang, 2022). Symmetry guesses were validated by initial helical refinement in cryoSPARC and selection of the helical symmetry parameters yielding reconstructions with typical PspA features and the best resolution. Then all segments were classified by heterogeneous refinement and subsequent 3D classifications using the initial helical reconstructions as templates. The resulting class distribution gave the PspA rod diameter distribution shown in Fig. 2 and Suppl. Fig. 2. The resulting helical reconstructions were subjected to multiple rounds of helical refinement including the symmetry search option. For the final polishing, the segments were reextracted at 400 px without Fourier cropping. Bad segments were discarded by heterogeneous refinement. Higher-order aberrations were corrected by global and local CTF refinement followed by a final helical refinement step. The local resolution distribution and local filtering was performed using cryoSPARC. The resolution of the final reconstructions was determined by Fourier shell correlation (auto-masked, FSC=0.143). The peaks in some FSC curves at 1/5.3 Å and lower resolution (Suppl. Fig. 2 and 6) correspond to the features (α-helix pitch) of all α-helical structures present in the PspA protein.

### Cryo-EM map interpretation and model building

The 3D reconstructions were B-factor sharpened in Phenix (*phenix.auto-sharpen*) (Terwilliger et al., 2018). The handedness of the final map was determined by rigid-body fitting the PspA reference structure aa 22-217 (PDB:7ABK) (Junglas et al., 2021) into the final maps using ChimeraX (Goddard et al., 2018; Pettersen et al., 2021) and flipped accordingly. 7ABK was flexibly MDFF fitted to the 3D reconstructions using ISOLDE (Croll, 2018). Then the respective helical symmetry was applied to all models to create assemblies of 60 monomers each. The assembly models were subjected to auto-refinement with *phenix.real_space_refine* (Afonine et al., 2018b) (with NCS constraints and NCS refinement). After auto-refinement, the new models were used for local model-based map sharpening with LocSCALE (Jakobi et al., 2017) to produce the final maps. The auto-refined models were checked/adjusted manually in Coot (Emsley et al., 2010) and ISOLDE (Croll, 2018) before a final cycle of auto-refinement with *phenix.real_space_refine* (Afonine et al., 2018b) (with NCS constraints and NCS refinement). After the final inspection, the model was validated in *phenix.validation_cryoem (Afonine et al., 2018a)/Molprobity (Williams et al., 2018).* Image processing, helical reconstruction, and model building were completed using SBGrid-supported applications (Morin et al., 2013). In this manner, from a total of 27 cryo-EM maps determined in three samples (apo, ADP, ATP), a total of unique 11 PDB coordinates were refined and deposited originating from the ATP sample (see **Table 1**). The remaining 16 maps were of identical symmetry consisting of either similar or poorer resolution densities and, therefore, atomic model refinement was not pursued further.

**Table 1.**
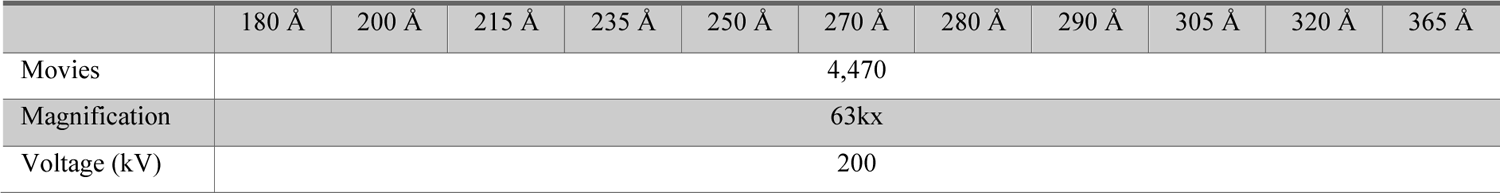
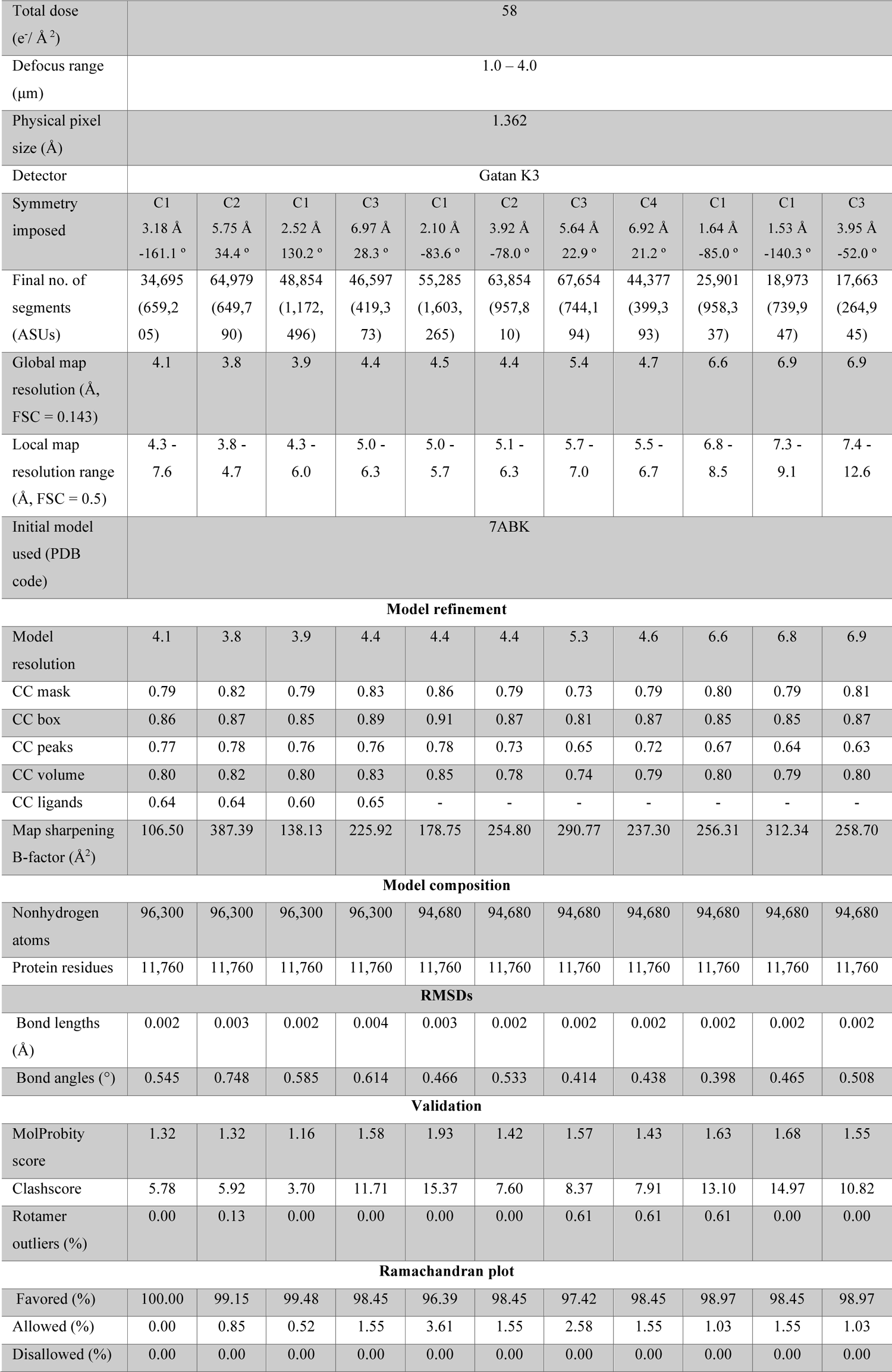
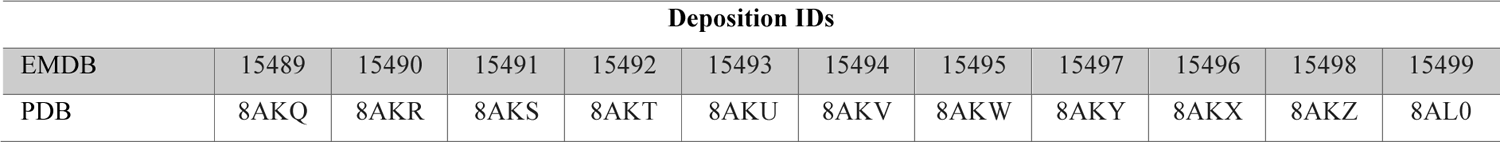
Data collection, image processing, and model refinement sample PspA+ATP.

Diameters of the final PspA rod reconstructions were measured using ImageJ (Rueden et al., 2017): First radial intensity profiles of each reconstruction were created. Then the radius at an intensity cutoff of 0.3 was read. The radius readout from reconstructions from the same helical symmetry class was averaged, converted to diameters, and rounded to 5 Å increments. Outer and inner leaflet radii of engulfed membrane tubes were determined from the same radial intensity profiles at the peak maxima from each bilayer leaflet.

### Analysis of membrane images

For description of the membrane morphology, subsets of 50 to 100 micrographs from sole SUVs (control), or SUVs with PspA or PspA+ATP, respectively, data sets were analyzed. Statistical analysis of the membrane features was performed on segmentations of the micrographs. Multiple micrographs from each dataset were manually segmented at a pixel size of 7 Å and given as patches as a training dataset to a standard U-Net (Ronneberger et al., 2015) with a depth of 4, patch size of 256, kernel size of 3 and batch size of 32. The micrographs were divided into patches, normalized to a mean of 0 and a standard deviation of 1, and simple rotations (90°, 180°, 270°) as well as flipped patches were added as data augmentation. The individual membranes are represented as the skeleton of their segmentation (Zhang and Suen, 1984), and highly aggregated or overlapping sections of the images were discarded as the automatic identification of the membrane shapes was ambiguous. Furthermore, only closed-segmented vesicles were analyzed. The perimeter of a vesicle was estimated by calculating the sum of the distances between neighboring points of the skeleton. For the automatic analysis of the enclosure distance, the skeleton points of a vesicle were compared with the points of other vesicles in the same micrograph. When the points were found within the skeleton of another vesicle, it was labeled as an enclosed vesicle. A vesicle can be enclosed by and enclose multiple other vesicles. When a vesicle was labeled as enclosed, the minimum Euclidean distance between the two vesicles was calculated. The thickness estimation of membranes was modified from Heberle and colleagues *(Heberle et al., 2022)*. Briefly, the image intensities are radially averaged along the membrane skeleton. The bilayer thickness for a membrane segment is then estimated by the distance between the minima in the radially averaged intensity profile for all nearby coordinates.

### Quantification and statistical analysis

Data and statistical analysis were performed using OriginPro 2021b (OriginLab Corp., Northampton, USA). Detailed descriptions of quantifications and statistical analyses (exact values of *n*, dispersion and precision measures used, and statistical tests used) can be found in the respective Figures, Figure legends, and Methods section.

## Author contributions

B.J., D.S., and C.S. designed the research. E.H., I.R., and A.H. cloned, expressed, and purified the proteins. B.J. and E.H. prepared cryo-EM samples. B.J. and E.H. operated the electron microscopes. B.J., E.H., and C.S. determined the cryo-EM structures. B.J. and E.H. built the refined atomic model. B.J., E.H., and A.H. measured and B.J., E.H., A.H., and D.S. interpreted the ATPase activity data. B.S.S. performed and B.S.S. and P.H. interpreted the mass spectrometry experiments. B.J., E.H., D.S., and C.S. prepared the manuscript with input from all authors.

## Declaration of interests

The authors declare no competing interests.

## Materials availability

All unique and stable reagents generated in this study are available from the Lead Contact with a completed Material Transfer Agreement.

## Data and code availability

The EMDB accession numbers for cryo-EM maps and PspA models are EMD IDs: 15489, 15490, 15491, 15492, 15493, 15494, 15495, 15497, 15496, 15498, 15499 and PDB-IDs: 8AKQ, 8AKR, 8AKS, 8AKT, 8AKU, 8AKV, 8AKW, 8AKY, 8AKX, 8AKZ, 8AL0.

## Acknowledgments

This study was funded by the Deutsche Forschungsgemeinschaft (DFG, German Research Foundation, SA 1882/6-1, SCHN 690/16-1, CRC1208 Project Nr 267205415 and CRC1551 Project Nr 464588647). The authors gratefully acknowledge the electron microscopy access time and computing time granted by the biological EM facility of the Ernst-Ruska Centre at Forschungszentrum Jülich. In this regard, we would thank Thomas Heidler and Pia Sundermeyer for maintaining the electron microscopes and Daniel Mann for maintaining the processing computers. The authors gratefully acknowledge the computing time granted through the Jülich Aachen Research Alliance (JARA) on the supercomputer JURECA at Forschungszentrum Jülich (Krause, 2019).

## Supplementary Information

**Suppl. Figure 1:**
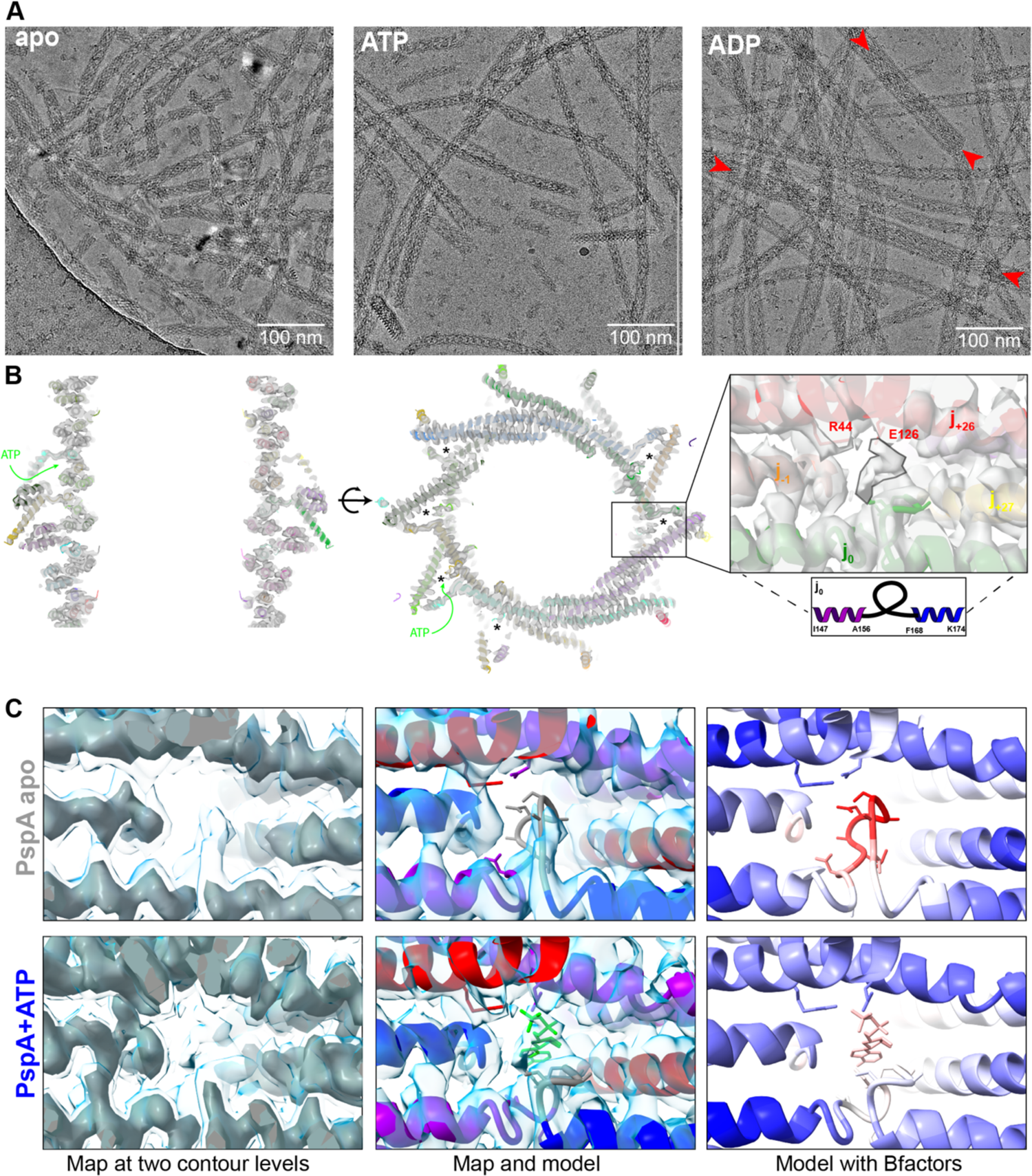
Example micrographs of PspA samples. A: Cryo-EM micrographs of the different PspA data sets. *Apo*: *PspA* without additives. ATP: PspA with 2 mM ATP. ADP: PspA with 2 mM ADP; red arrows show “super rods” with engulfed “normal” rods. **B:** Sliced side and top view of the PspA cryo-EM density in the presence of ATP (200 Å diameter, PDB: 8AKR, EMDB: 15490) with a superimposed atomic PspA model. Putative nucleotide-binding sites are labeled with asterisks. Green arrows indicate solvent accessibility of the ATP binding site. Inset: Zoomed view of the putative nucleotide-binding site in the PspA rod structure (200 Å diameter) with weak segmented density above the loop connecting helices α3 and α4 (cyan) due to incomplete binding and imposition of helical symmetry. The nucleotide-binding site is formed by the surrounding chains (subunits j0, j-1, j+26, and j+27) and is repeated for every subunit in the polymer. Inset. View of the cryo-EM density of the nucleotide binding region superimposed with PspA, no nucleotide model included in density (black rim). **C**: Comparison of the PspA apo structure with PspA after incubation with ATP. Left column: Maps of PspA and PspA+ATP at low and high contour levels superimposed. Middle column: Models of PspA apo (PDB:7ABK) and PspA+ATP with an ADP model fitted to the additional density. Note that due to the presence of the nucleotide the loop in the PspA+ATP structure is moved. Right column: Atomic models of PspA apo and PspA+ATP superimposed with colored atomic temperature factors indicate increased mobility of α3/α4 loop in the PspA apo structure.

**Suppl. Figure 2:**
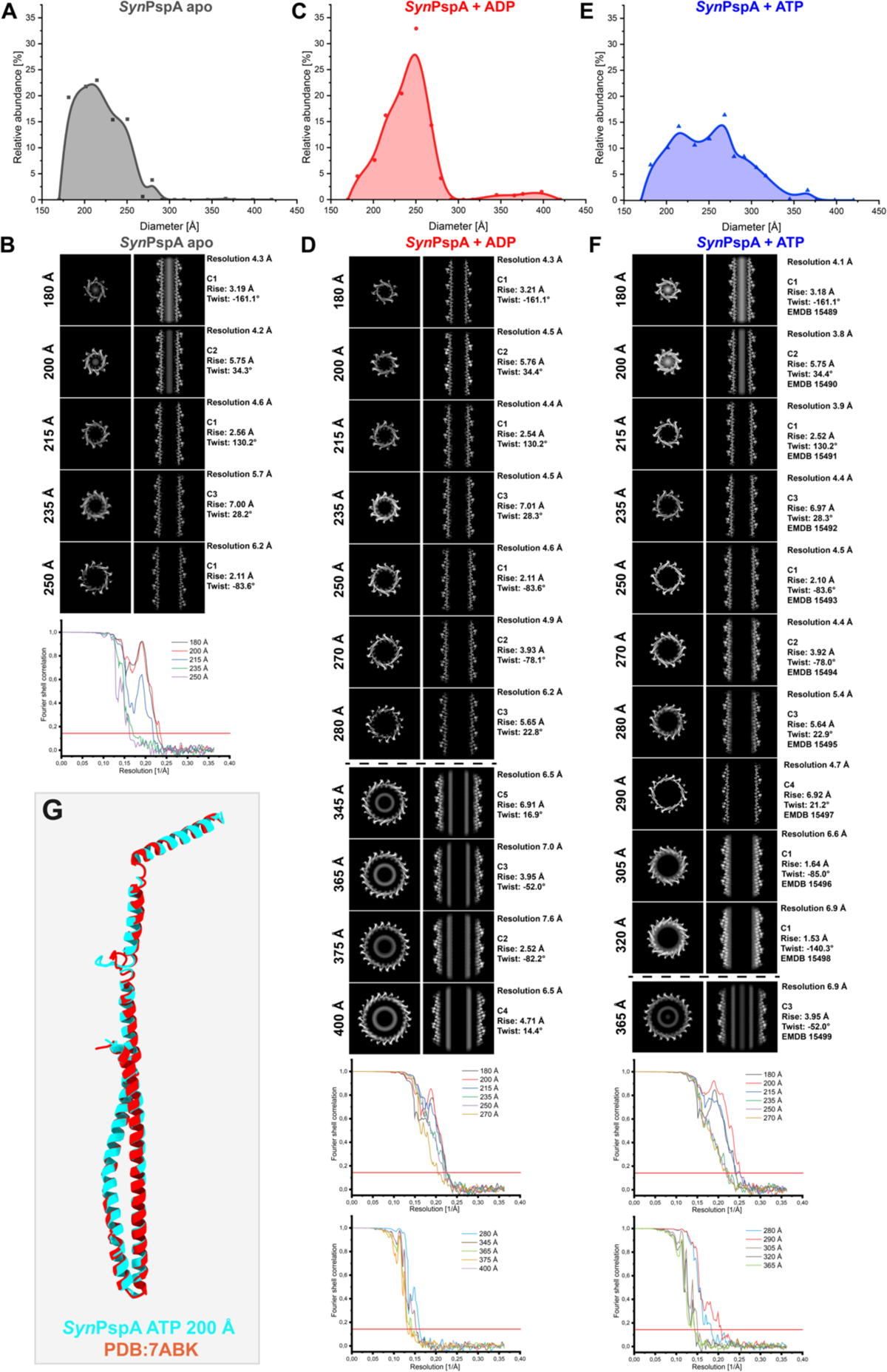
PspA rod diameter distributions and corresponding cryo-EM structures in the absence or presence of nucleotides. PspA rod diameter distribution for **A:** PspA *apo* (grey), **C:** PspA+ADP (red), and **E:** PspA+ATP (blue), based on the relative occurrence of rod segments with a certain diameter. **B, D**, **F:** Overview of PspA rod cryo-EM structures with cross-sectional top view *z*-slices (left column), cross-sectional side view *xy*-slices (middle column) and FSC curves with a 0.143 threshold of **B:** PspA *apo*, **D:** PspA+ADP and **F:** PspA+ATP. Note that when FSC curves drop below 1, systematic peaks of high correlation occur at 1/5.3 Å corresponding to the α-helical pitch feature and additional low-resolution details of the all α-helical maps of PspA. **G:** Comparison of the PspA ATP 200 Å monomer with the high-resolution reference structure of PspA *apo* PDB:7ABK.

**Suppl. Figure 3:**
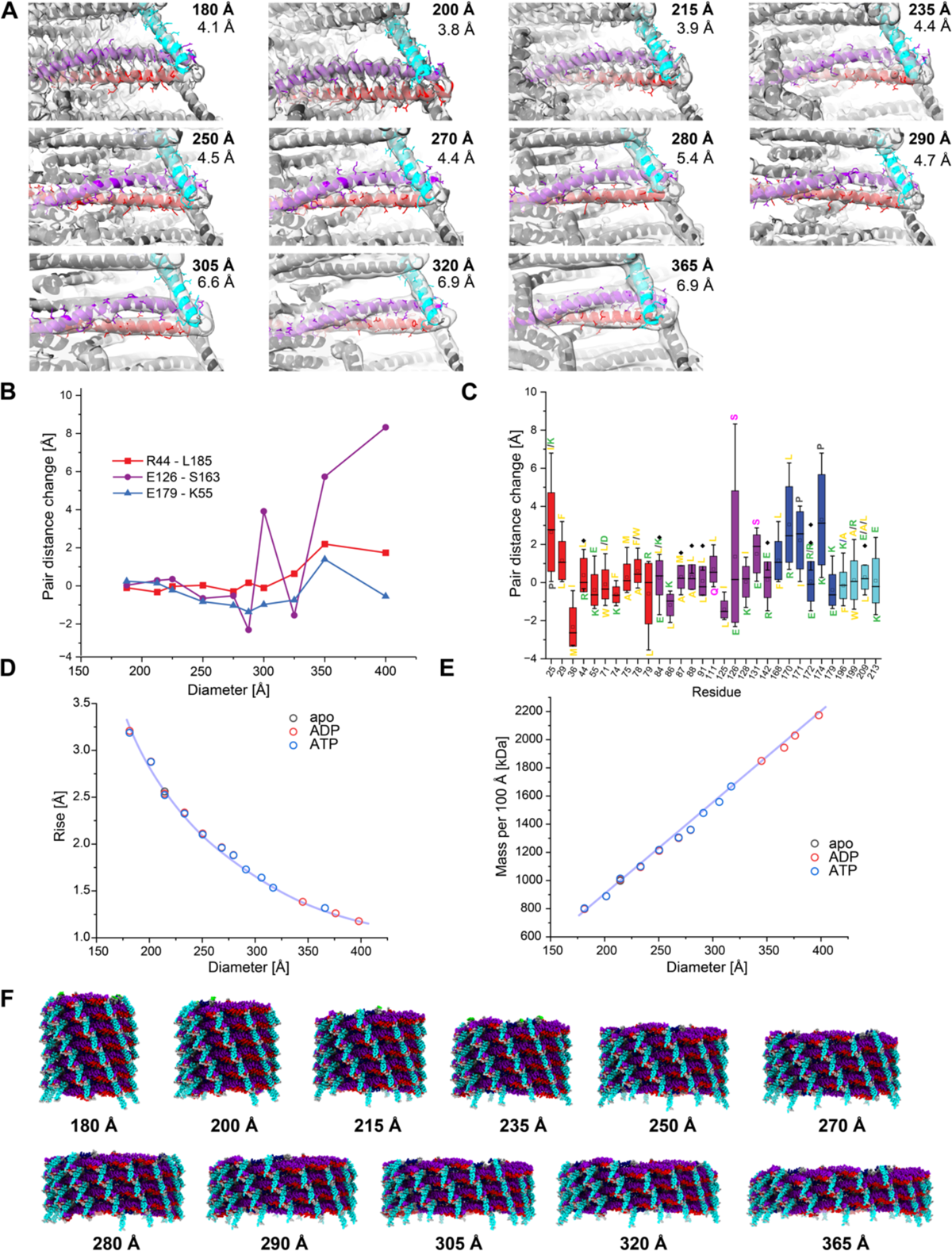
PspA rod radial density profiles and mass per length. A: Cryo-EM density fit of atomic PspA models of different diameters focusing on the helices α1/α2 hairpin and helix 5 ((1 red, (2+3 violet, (4 blue, (5 cyan). (180 Å: PDB 8AKQ, 200 Å: PDB 8AKR, 215 Å: PDB 8AKS, 235 Å: PDB 8AKT, 250 Å: PDB 8AKU, 270 Å: PDB 8AKV, 280 Å: PDB 8AKW, 290 Å: PDB 8AKY, 305 Å: PDB 8AKX, 320 Å: PDB 8AKZ, 365 Å: PDB 8AL0) B: Scatter plot of three selected pairs (R44-L185, E126-S163, E179-K55) distance changes with respect to the initial distance in the 180 Å diameter assembly over all rod diameters. C: Box plot of pair distance changes of evolutionarily conserved residues with respect to the initial distance in the 180 Å diameter assembly. The residues were selected by first identifying potential intermolecular interactions between highly conserved residues (>90% conserved among PspA/Vipp1proteins (Junglas, et al., 2021). Then, the C( distance for each pair was measured for each rod diameter. To calculate the difference of the pair distances relative to the smallest diameter rods, the distances in 180 Å rods were subtracted from the respective distances in the other diameters. The standard deviation (SD) of the distance shift distribution is a measure of the flexibility of the interaction. Boxes show SD with median line (line) and mean value (circle). Whiskers show the range within 1.5 IQR. Color code for residues: green=charged; yellow=hydrophobic; pink=polar+uncharged: grey=special cases. Color code for boxes: residues located in helix α1: red, residues located in helix α2+3: violet, residues located in helix α4: blue, residues located in helix α5: cyan. D: Diameters plotted against helical rise (after correction for number of strands) in Å. *Apo* (black): PspA ADP (red): PspA + 2mM ADP. ATP (blue): *PspA* + 2 mM ATP. E: Diameters plotted against mass per 100 Å length in kDa. *Apo* (black): *PspA*. ADP (red): *PspA* + 2mM ADP. ATP (blue): PspA + 2 mM ATP. F: PspA models with 60 monomers each show a decrease in rod length and an increasing diameter from 180 - 365 Å.

**Suppl. Figure 4:**
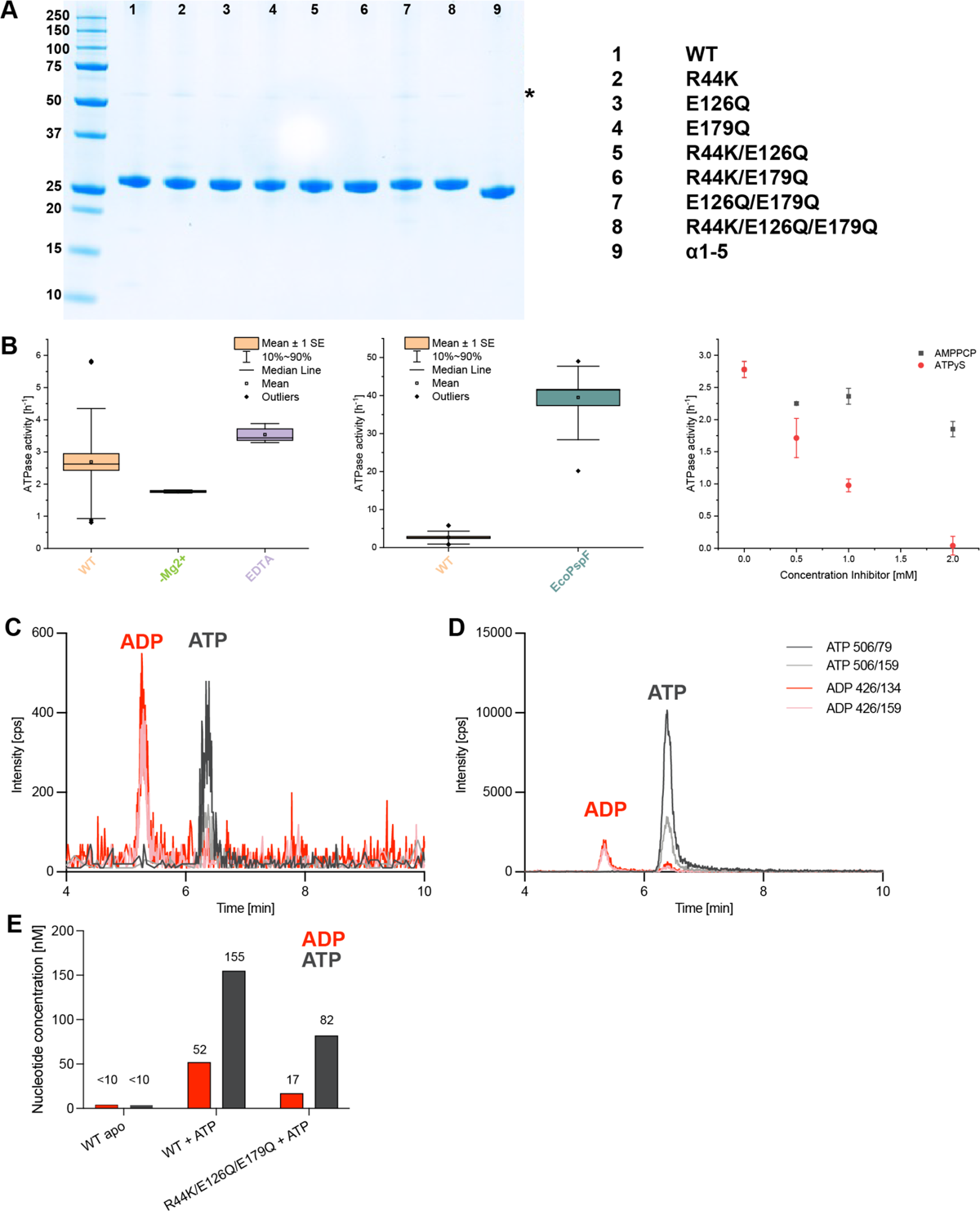
Nucleotide binding and hydrolysis by PspA wild-type (WT) and mutants. **A:** SDS PAGE of purified PspA WT and mutants (3 µg each). Purified proteins show a band at 28 kDa, except for the α0 truncated form which shows a band at 25 kDa. Asterisk: Faint dimer bands at twice the height of the monomeric protein band (56 kDa for WT and full-length PspA mutants; 50 kDa for α1-5 mutant). Marker: Bio-Rad Precision PlusProteinTM Unstained standards. **B:** ATPase activity of PspA. Left graph: Influence of Mg^2+^ and EDTA on the ATPase activity of the WT protein. WT (orange): ATPase activity in the presence of 2 mM Mg^2+^. -Mg^2+^(green): ATPase activity in the absence of Mg^2+^. EDTA (purple): ATPase activity in the presence of 10 mM EDTA; n(WT)=36, n(-Mg^2+^ and EDTA)=3. The boxplots show the mean ± standard error as boxes, the 10% to 90% percentile as whiskers, and outliers as diamonds. Middle graph: Comparison of the PspA ATPase activity with ATPase activity of *E. coli* PspF, n(PspA)=36, n(*Eco*PspF)=3. The boxplots show the mean ± standard error as boxes, the 10% to 90% percentile as whiskers, and outliers as diamonds. Right graph: Influence of ATPyS and AMPPCP on the ATPase activity of PspA. Red circle: ATPyS; Grey square: AMPPCP; n=3, error bars represent SD. **C:** HPLC/MS-MS of PspA directly after purification. **D:** HPLC/MS-MS of PspA R44K/E126Q/E179Q + ATP after 24 h incubation and extensive washing. Different color lines represent different MRM transitions. For ADP, MRM transitions are 426/134 (red) and 426/159 (light red). For ATP MRM transitions are 506/79 (dark grey) and 506/159 (light grey). The ADP fragments below the ATP peak are formed by in-source fragmentation of ATP in the ESI source. **E:** Bar plot of estimated nucleotide concentrations found by LC-MS/MS in PspA *apo* control, WT, and R44K/E126Q/E179Q rods after incubation with ATP for 24 h and extensive washing, n=1.

**Suppl. Figure 5:**
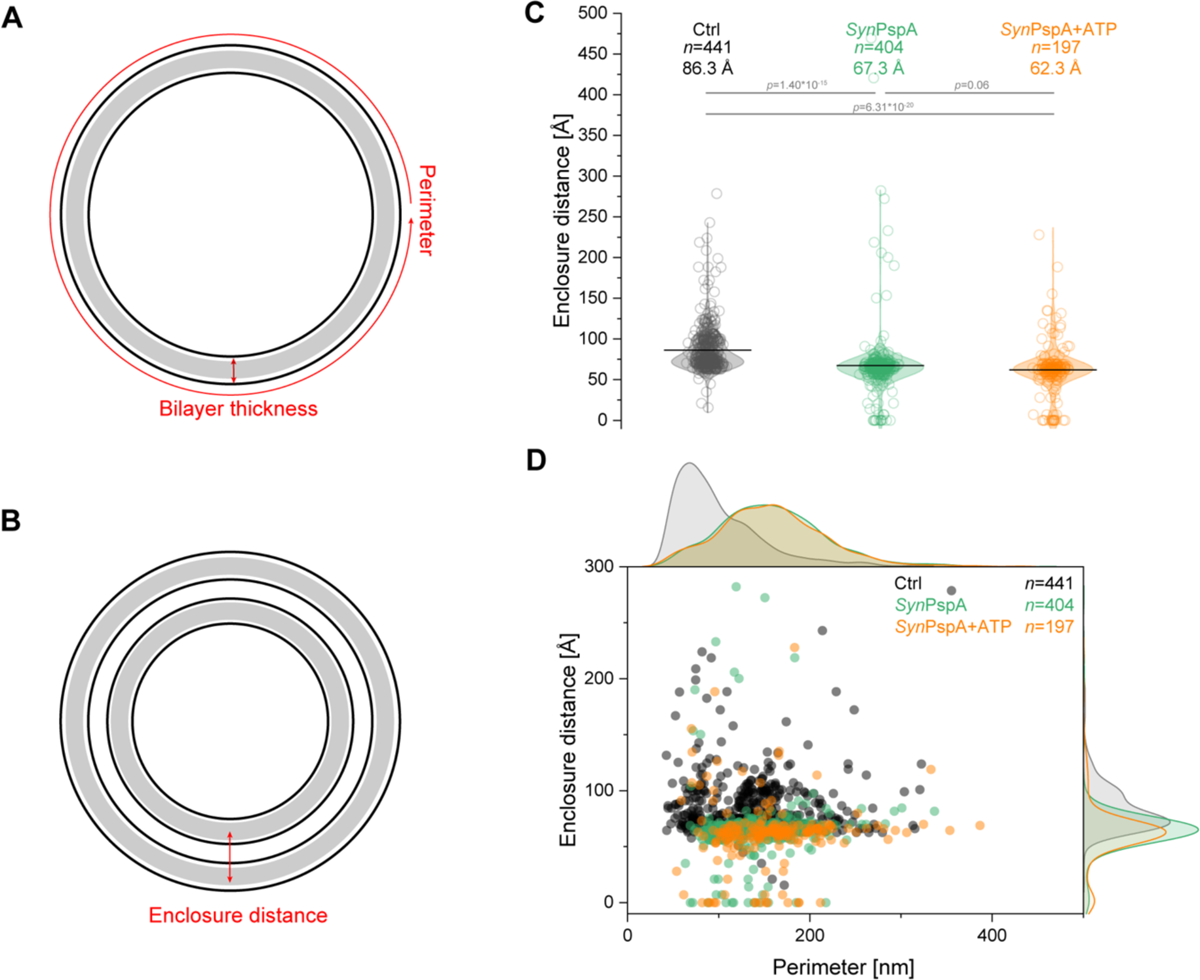
Morphological changes of EPL vesicles after *Syn*PspA/ATP reconstitution. A+B: Schematic view of the measured vesicle parameters: perimeter, bilayer thickness, and enclosure distance (see materials and methods for details). C: Violin plot of the enclosure distance of control EPL SUVs (grey), PspA+EPL SUVs (green), and PspA+EPL+ATP SUVs (orange). The mean enclosure distance, and the number of measured vesicles *n* and *p* values from a two-sample t-test are indicated on the graph. D: Scatter plot and histogram of the vesicle perimeter versus enclosure distance of control EPL SUVs (grey), PspA+EPL (green), and PspA+EPL+ATP (orange). The number of measured vesicles *n* is indicated on the graph.

**Suppl. Figure 6:**
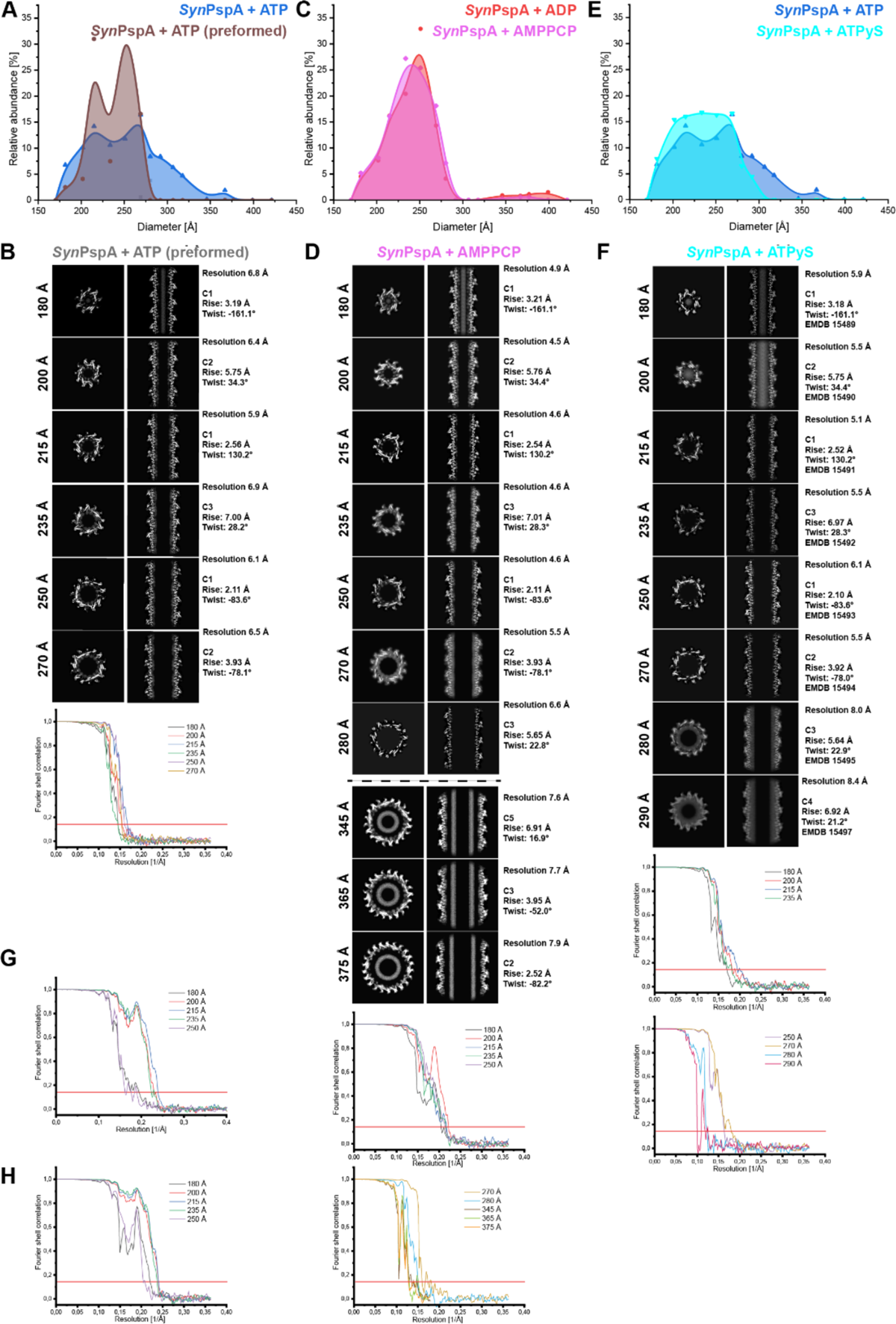
PspA rod diameter distribution and corresponding cryo-EM structures in the presence of non-hydrolyzable ATP analogs. **Top.** PspA rod diameter distribution for **A:** PspA+ATP (blue), and preformed PspA rods incubated with ATP (brown), **C:** PspA+ADP (red) and PspA+AMPPCP (magenta), and **E:** PspA+ATP (blue) and PspA+ATPyS (cyan), based on the relative occurrence of rod segments with a certain diameter. **Bottom.** Cross-sectional top view *z*-slices (left column), cross-sectional side view *xy*-slices (middle column) and FSC curves (threshold 0.143) of cryo-EM structures **B:** PspA+ATP (preformed), **D:** PspA+AMPPCP and **F:** PspA+ATPyS. **G:** FSC curve of PspA dL10 diameters (threshold 0.143). **H:** FSC curve of PspA dL10 diameters with ATP (threshold 0.143). Note that when FSC curves drop below 1, systematic peaks of high correlation occur at 1/5.3 Å corresponding to the α-helical pitch and additional low-resolution features of the all α-helical maps of PspA.

---

**Suppl. Table 1:** Parameters for the quantification of ATP/ADP.

| Compound | Precursor<br>(*m/z*) | Product ion<br>(*m/z*) | DP<br>(*eV*) | CE<br>(*eV*) | CXP<br>(*eV*) | Retention<br>time (min) |
| --- | --- | --- | --- | --- | --- | --- |
| ATP | 506 | 79 (quantifier) | -105 | -30 | -11 | 6.3 |
|  |  | 159 (qualifier) | -105 | -34 | -9 |  |
| ADP | 426 | 134 (quantifier) | -120 | -39 | -7 | 5.2 |
|  |  | 159 (qualifier) | -120 | -34 | -9 |  |

**Suppl. Table 2.** Sample Details.

|  | *SynPspA*<br>*apo* | *SynPspA*<br>*ATP* | *SynPspA*<br>*ADP* | *SynPspA*<br>*AMPPC*<br>*P* | *SynPspA*<br>*ATPyS* | *SynPspA*<br>*dL10* | *SynPspA*<br>*dL10*<br>*ATP* | *SynPspA*<br>*ATP*<br>*preformed* | *SynPsp*<br>*A EPL* | *SynPsp*<br>*A EPL*<br>*ATP* | *SynPspA*<br>*R44K/E1*<br>*26Q/E17*<br>*9Q/ EPL*<br>*ATP* | *EPL* |
| --- | --- | --- | --- | --- | --- | --- | --- | --- | --- | --- | --- | --- |
| Protein<br>Conc. | 6 – 8<br>mg/m<br>L | 6 – 8<br>mg/m<br>L | 6 – 8<br>mg/m<br>L | 6 – 8<br>mg/m<br>L | 6 – 8<br>mg/m<br>L | 0.5<br>mg/m<br>L | 0.5<br>mg/m<br>L | 0.1<br>mg/mL | 1.2<br>mg/m<br>L | 1.2<br>mg/m<br>L | 0.9<br>mg/mL | - |
| Lipid<br>Conc. | - | - | - | - | - | - | - | - | 2.5<br>mg/m<br>L | 2.5<br>mg/m<br>L | 2.5<br>mg/mL | 2.5<br>mg/m<br>L |
| NTP<br>Conc. | - | 2 mM | 2 mM | 1 mM | 0.5 | - | 2 mM | 2 mM | - | 10<br>mM | 10 mM | - |
| MgCl<sub>2</sub><br>Conc. | - | 4 mM | 4 mM | 2 mM | 4 mM | - | 4 mM | 4 mM | - | 20<br>mM | 20 mM | - |
| Magnifi<br>cation | 63 kx | 63 kx | 63 kx | 63 kx | 63 kx | 105 kx | 63 kx | 63 kx | 63 kx | 63 kx | 63 kx | 63 kx |
| Pixel<br>size | 1.362<br>Å | 1.362<br>Å | 1.362<br>Å | 1.362<br>Å | 1.362<br>Å | 0.84 | 1.362<br>Å | 1.362 Å | 1.362<br>Å | 1.362<br>Å | 1.362<br>Å | 1.362<br>Å |
| Frames | 40 | 40 | 40 | 40 | 40 | 35 | 45 | 45 | 40 | 40 | 40 | 40 |
| Total<br>dose | 58<br>e-/Å<sup>2</sup> | 58<br>e-/Å<sup>2</sup> | 58<br>e-/Å<sup>2</sup> | 58<br>e-/Å<sup>2</sup> | 58<br>e-/Å<sup>2</sup> | 35<br>e-/Å<sup>2</sup> | 45<br>e-/Å<sup>2</sup> | 45<br>e-/Å<sup>2</sup> | 58<br>e-/Å<sup>2</sup> | 58<br>e-/Å<sup>2</sup> | 58<br>e-/Å<sup>2</sup> | 58<br>e-/Å<sup>2</sup> |
| Defocus<br>range | 1.0 to -<br>3.0 µm | 1.0 to<br>4.0 µm | 1.5 to<br>3.5 µm | 1.5 to<br>3.5 µm | 1.0 to<br>4.0 µm | 1.5 to<br>3.0 µm | 1.5 to<br>3.0 µm | 1.5 to<br>3.0 µm | 1.5 to<br>3.5<br>µm | 2.0 to<br>4.0<br>µm | 2.0 to<br>4.0 µm | 2.0 to<br>4.0<br>µm |
| Movies | 1,818 | 4,470 | 3,038 | 3,127 | 3,310 | 3,620 | 4,311 | 3,065 | 50 | 61 | 73 | 632 |

## Notes

### Competing Interest Statement

The authors have declared no competing interest.

